# Pathway specific activation of striatal APJ receptor suppresses dopamine D1 receptor sensitization and L-DOPA induced dyskinesia

**DOI:** 10.64898/2026.11.16.766711

**Authors:** Mohamed Rafiuddin Ahmed, Chen Zheng, Mohamed Sohail Ahmed, Mithya Morton, Mustafeez Mujtaba Babar, Mohammed Inayathullah, Jayakumar Rajadas

## Abstract

L-DOPA induced dyskinesia (LID) remains a major debilitating complication in the long-term management of Parkinson’s disease (PD), primarily driven by the hypersensitization of dopamine D_1_ receptors (D_1_R) on striatal direct pathway medium spiny neurons (MSNs). In this study, we comprehensively investigated the therapeutic efficacy, translational feasibility, cellular mechanisms, and circuit-level logic of apelin peptide administration targeting the APJ receptor (APJR) in hemiparkinsonian mouse models. Utilizing adeno-associated virus (AAV) gene delivery, non-invasive liposomal intranasal delivery, and cre dependent cell type specific viral manipulations, we evaluated behavioral sensitization, abnormal involuntary movements (AIMs), and intracellular signaling dynamics. Striatal AAV-mediated expression of apelin-36 attenuated L-DOPA-induced rotational sensitization in hemiparkinsonian mice. From a translational standpoint, non-invasive intranasal delivery of liposomal apelin13 markedly reduced AIMs, while completely preventing progressive rotational sensitization over chronic L-DOPA treatment regimens. Mechanistically, apelin peptide treatment attenuated the persistent nuclear accumulation of the dyskinesia transcription factor marker ΔFosB in lesioned striatal tissue and normalized arrestin3 isoform expression. Although the ERK1/2 and p38 mitogen-activated protein kinase (MAPK) pathways were hyperactivated in the lesioned striatum, there was no difference between control and apelin peptide treated animals, however we observed reduced downstream phospho-mTOR levels with apelin13 peptide treatment. Cre-dependent expression of apelin-36 in Drd1a-expressing, but not Adora2a-expressing, neurons reduced L-DOPA-induced rotational sensitization. Together, these findings demonstrate that targeting the striatal APJR axis on direct pathway neurons provides a potent, non-dopaminergic therapeutic strategy to counteract D1R hyper-sensitization and alleviate LID in PD.

## Introduction

Parkinson’s disease (PD) is a debilitating neurodegenerative disorder primarily characterized by motor symptoms such as bradykinesia, rigidity, tremor, and postural instability^1,2^. The disease results from the progressive loss of dopaminergic neurons in the substantia nigra pars compacta, leading to a significant reduction in dopamine (DA) levels within the striatum^2–5^. The gold standard treatment for PD involves DA replacement therapy using L-3,4-dihydroxyphenylalanine (L-DOPA), which is converted to DA in the brain to alleviate motor symptoms^6^. Despite its efficacy, chronic L-DOPA treatment often leads to a severe complication known as L-DOPA induced dyskinesia (LID)^7,8^. LID is characterized by abnormal, involuntary movements that can be as debilitating as the primary symptoms of PD^9^. The development of LID is associated with the sensitization and aberrant signaling of DA receptors, particularly within the striatum^10,11^. However, the precise molecular mechanisms underlying these maladaptive changes remain elusive and poorly understood^12–15^. This gap in knowledge has hindered the development of effective strategies to prevent or treat LID, necessitating further investigation into the molecular events and signaling pathways involved. The DA receptors belonging to the GPCR family, play a crucial role in modulating motor function and other neurological adaptations in the brain^14^. Chronic L-DOPA administration under DA depleted conditions such as PD, alters DA receptor (DAR) signaling contributing to plasticity and pathophysiology of LID^16–18^. Studies have shown that LID is associated with increased DAR sensitivity, changes in receptor phosphorylation, and altered intracellular signaling pathways^11,19–22^. These studies have predominantly focused on DA receptors, with limited exploration of other interacting partners with DARs that may modulate intracellular signaling events and may consequentially contribute or ameliorate LID.

It is a well-established paradigm that in the striatum, long-term administration of L-DOPA in PD animal models leads to maladaptive synaptic plasticity and LID^13^. Although hypersensitized DA D_1_ receptor (D_1_R) signaling in direct pathway spiny projection neurons (dSPNs) drives this abnormal response, non-dopaminergic G protein coupled receptors (GPCRs) critically modulate these circuits and influence dyskinesia severity^23^. Postsynaptic adenosine A_2A_ receptors (A_2AR_), selectively expressed on indirect pathway spiny projection neurons (iSPNs), form functional heteromeric complexes with DA D_2_ receptors (D_2_R), where A_2A_R antagonism enhances D_2_R signaling to tone down indirect pathway overactivity^24,25^. Furthermore, presynaptic A_2A_R and metabotropic glutamate receptor 5 (mGluR_5_) fine tune corticostriatal glutamatergic drive onto SPNs and targeting these receptors with A_2A_R antagonists or mGluR_5_ negative allosteric modulators reduces excessive striatal excitation and ameliorates dyskinetic movements^25,26^. Central serotonin 5HT_1A_ and 5HT_1B_ auto receptors on serotonergic terminals control unbuffered, non-physiological DA release from converted L-DOPA, while postsynaptic 5HT receptors on SPNs crosstalk with dopaminergic cascades to recalibrate basal ganglia output^13^.

Crucial emerging evidence implicates that apelin receptor (APJR), a dual G_i/o_ and G_q_ coupled GPCR abundantly expressed in striatal SPNs, as a powerful non-dopaminergic regulator of direct pathway transmission^27–32^. APJR forms functional heteromeric complexes with other GPCRs within SPNs^33–37^. Through physical protomer crosstalk, APJR activation could potentially exert an allosteric trans-inhibition of other GPCR signaling^38–41^. We hypothesized that co-activation or biased pharmacological targeting of APJR could dampen DAR especially, D_1_R stimulated G_αs/olf_ coupling and adenylyl cyclase (AC) activity preventing the pathological surge in intracellular cAMP accumulation and downstream protein kinase A (PKA)/DARPP-32 hyperphosphorylation that characterizes LID. Moreover, ligand-induced engagement of APJR could shift D_1_R dynamic signaling toward preferential β-arrestin recruitment and GRK-mediated receptor desensitization. This biased signaling profile effectively uncouples D_1_R from dyskinesiogenic canonical cAMP/PKA cascades while promoting receptor endocytosis and recalibrating dSPN excitability. Together, these non-dopaminergic GPCR systems shape SPN excitability, intracellular signaling cascades, and receptor crosstalk, highlighting their relevance as therapeutic targets to mitigate LID without compromising the motor benefits of L-DOPA.

## Materials and methods

### Plasmids for transfection

The pcDNA3 human DAR-3X HA tag (D1R-3X-HA and D2R-3X-HA long isoform), pcDNA3-hD1R-LgBiT, muscarinic 2 receptor (3X-HA-M2R-LgBiT), arrestin2-SmBiT, arrestin3-SmBiT, arrestin2-GFP and arrestin3-GFP were a kind gift from Dr. Seva Gurevich lab (Vanderbilt University). The APJR-DuET plasmid was from Dr. Thomas Sakmar (The Rockefeller University) and was obtained from Addgene (Plasmid#213185)^42^. The APJR-DuET plasmid was reengineered to remove the DuET sequence and inserted with the LgBiT sequence for glosensor cAMP accumulation assay using standard PCR methods and sequenced by next generation sequencing.

### NanoBiT arrestin recruitment assay

HEK293 arrestin-2/3 double knockout (dko) cells were seeded into dishes and transfected using TransHi transfection reagent (FormuMax, Sunnyvale CA) upon reaching 80-90% confluence using a vector combination consisting of GPCR-LgBiT, SmBiT-arrestin2 or arrestin3, and pcDNA3 carrier plasmid. Twenty-four hours post-transfection, cells were detached using 0.05% trypsin-EDTA, resuspended in phenol red free DMEM (GIBCO) containing 10% FBS, and harvested by centrifugation at 400 × *g* for 5 min. Cell pellets were resuspended in serum free, phenol red free DMEM, seeded into white flat bottom 96 well plates at 100 µL per well, and incubated overnight at 37 °C for cell starvation. On the following day, 20 µL of coelenterazine h substrate (diluted 1:50 in DPBS) were added to each well, and baseline luminescence was recorded every minute for 10 min on a microplate reader to establish a stable signal. Cells were then stimulated by adding 5 µL of agonist solution, and luminescence kinetics were monitored continuously for 30-60 min as described^43^.

### Real-time glosensor cAMP accumulation assay

HEK293 cells were seeded into white flat-bottom 96-well plates and grown overnight to achieve 90-95% confluence. Cells were transfected with equal amounts (50 ng per well) of receptor plasmid DNA and Glosensor 22F cAMP reporter construct (Cat# E1291, Promega Corporation, Madison WI) using TransHi transfection reagent in serum-free medium. At 24 h post-transfection, the culture medium was exchanged for serum free DMEM. At 48 h post-transfection, cells were incubated for 2 h at 37 °C with Glosensor cAMP reagent (2% v/v in serum-free medium). Following incubation, baseline luminescence was recorded for 20 min to assess background signal. Target ligands (10µL diluted in serum-free medium) were then added to achieve a final reaction volume of 100µL, and real-time luminescence emission was continuously recorded for 2h. Data were collected and analyzed using GraphPad Prism software.

### Liposomal drug formulations

The apelin13 liposomal formulations were made using DSPC and DPPC at a 1:1 ratio (80% w/w), as described^44^. The liposomes were further stabilized by adding polyethylene glycol (PEG) 8000 at 8% (w/w) and cholesterol at 4% (w/w) to the final liposomal weight. The lipids were dissolved in chloroform and were then evaporated in vacuum using rotavapor. The thin film thus formed was reconstituted in DNAse/RNAse-free water. HPLC purified apelin13 peptide (CS Bio Palo Alto, CA) at a concentration of 20mg/mL was added to the reconstituted lipid cake. The reconstituted aqueous peptide liposomal complex was freeze-dried, reconstituted, and freeze-dried three times for maximal drug encapsulation.

### Immunocytochemistry and confocal microscopy

HEK293 wildtype and arrestin 2/3 dko cells were seeded onto poly-D-lysine coated glass coverslips placed inside cell culture plates and cultured overnight to allow proper attachment. Following transfection with the indicated receptor and reporter constructs, cells were subjected to specified ligand treatments or vehicle conditions. Cells were then washed with ice cold phosphate buffered saline and fixed using 4% paraformaldehyde in phosphate buffered saline (PBS) for 15 minutes at room temperature. After fixation, cells were washed three times with PBS and permeabilized with 0.1% Triton X 100 (in PBS) for 10 minutes. To prevent nonspecific antibody binding, cells were incubated in blocking buffer consisting of 5% bovine serum albumin in PBS for 1 hour at room temperature. Coverslips were then incubated overnight at 4 °C with primary antibodies targeted against the candidate receptors (HA tag or APJR ab), arrestin isoforms (GFP, mouse JL8 Clontech), or specific subcellular markers. After washing off unbound primary antibodies with PBS, cells were incubated for 1 hour at room temperature in the dark with fluorophore conjugated secondary antibodies. Nuclei were counterstained with DAPI for 5 minutes. Coverslips were mounted onto microscope slides using anti fade mounting medium and sealed. Fluorescent signals were visualized and imaged using a Nikon laser scanning confocal microscope equipped with appropriate laser lines and emission filters. Optical sections were acquired using a high numerical aperture oil immersion objective, and laser power and detector gain settings were maintained consistently across all experimental groups to enable accurate comparison of receptor localization, translocation, and arrestin dependent trafficking events between wild type and arrestin 2/3 dko cells.

### Adeno-associated virus construction and preparation

The apelin36 and scrambled apelin36 gene was custom synthesized by Blue Heron (Bothel, WA, USA) and was tagged with FLAG at the C-terminus. The viral construct was assembled in the adeno-associated viral vector (AAV) under control of the EF1α promoter and included downstream of it co-cistronic GFP under control of super-IRES (sIRES), which has been shown to significantly increase protein expression^45,46^. GFP alone was used as a control. The double inverted open reading frame (DIO) constructs for the expression of apelin36 and scrambled apelin36 peptides along with co-cistronic expression of GFP were amplified using PCR and subcloned into the AAV-CMV-CAG-DIO-Av vector (CMV with chicken actin growth factor) (a kind gift from Dr. Mark Schnitzer Lab, Stanford University) at PstI and AscI (5-3’) cloning sites flanked by the loxP sites for expression under cre. The adeno associated viruses serotype 5 (AAV-Rep/Cap-5) encoding GFP without the peptide sequence was used as a control. The AAV-apelin36/scrambled apelin36 (with co-cistronic GFP) under control of either EF1α or CAG-CMV promoter were constructed and produced using AAV Helper Free Expression System and 293-AAV cell line (Cell Biolabs, Inc.). The viral tier was determined by standard qPCR with the BioRad SYBR Green supermix using serial dilutions of the cloning vector pAAV-MCS as standards.

### Hemiparkinsonian mouse model

c57Bl/6 mice from Jackson laboratories, aged 12 to 20 weeks old, inbred dopamine D1cre and A_2A_-cre lines (shared from Dr. Mark Schnitzer’s laboratory, Stanford University and Dr. Gurevich laboratory, Vanderbilt University), were used for all hemiparkinsonian studies. Adult male and female mice were housed in a Stanford and Vanderbilt University animal facilities with a 12/12 hr light/dark cycle and had water and food ad libitum. All procedures followed National Institute of Health guidelines and were approved by the institutional animal care and use committee (IACUC) of both the institutes. Mice were deeply anesthetized with 5% isoflurane (Isothesia, Henry Schein Animal Health, Dublin, OH) in 100% oxygen with a delivery rate of 5 L/minute until loss of righting reflex and mounted on a stereotaxis frame as previously described^44,47^. The animals were maintained under anesthesia with 1-1.5% isoflurane throughout the surgical procedure. Body temperature was monitored throughout the surgical procedure; respiration was monitored every 10 minutes till the end of the procedure. Before the infusion of 6-hydroxydopamine (6-OHDA), mice were treated with desipramine HCl (25 mg/kg i/p) to protect the non-adrenergic neurons from damage. 2 μL (2 μg/μL) of 6-OHDA (Santacruz Biotechnology, Santa Cruz, CA) was unilaterally infused at the rate of 0.5 μL/min into the median forebrain bundle (MFB) at coordinates AP=1.1, ML=-1.3, DV=5.02 from the bregma to target nigral dopaminergic projection neurons. Animals were allowed to recover from anesthesia. Four weeks after lesion surgery, animals were tested for behavior.

### Behavioral assessments in the hemiparkinsonian mice model

Unilateral 6-OHDA lesioning of the medial forebrain bundle in mice creates a validated hemiparkinsonian model characterized by severe striatal dopamine depletion. To evaluate L-DOPA induced dyskinesia and motor asymmetry, mice were subjected to standardized abnormal involuntary movement scoring and rotational behavioral assessments following intranasal apelin peptide or vehicle treatments and by AAV mediated expression of apelin36 peptide.

### Abnormal involuntary movements (AIMs)

Abnormal involuntary movements were scored by an observer blinded to the experimental conditions to quantify the severity of dyskinesia. Individual mice were placed in clear plexiglas observation chambers and habituated for at least 30 minutes. The mice were administered via intranasal route the liposomal formulation of apelin13 peptide or blank formulations without any peptide 30 minutes before L-DOPA injection at 5mg/Kg body weight. The AIMs were evaluated at 20-minute intervals over 180-minute period. Dyskinesia was categorized into distinct subtypes: axial dystonia, characterized by twisted postures of the neck and torso toward the side contralateral to the lesion; limb dyskinesia, observed as repetitive purposeless movements or dystonic posturing of the contralateral forelimb; and orolingual dyskinesia, defined as vacuous chewing, tongue protrusions, and facial twitching. Each subtype was scored on a severity scale ranging from 0 to 4 based on its persistence during each observation window, where 0 indicated the absence of dyskinetic movements, 1 represented sporadic movements present for less than half the observation time, 2 indicated frequent movements present for more than half the time, 3 signified continuous movements interrupted only by external sensory stimuli, and 4 denoted continuous uninterrupted dyskinesia. Subtype scores were analyzed individually and summed to generate a total AIM score for each time point and session.

### Rotational behavior

Rotational behavior was measured to assess motor asymmetry and functional hypersensitivity of striatal dopamine receptors on the lesioned side. Mice were fitted with specialized harnesses and connected to automated rotometer plexiglas cylinders (Omnitech Instruments, Ohio) designed to record rotational turns continuously. The mice were administered via intranasal route similar to the AIM study with liposomal apelin13 peptide or blank formulations minus peptide and followed by L-DOPA injection at 25mg/Kg body weight. Following administration of L-DOPA, net rotations were recorded for up to 60 minutes. Contralateral rotations, defined as full 360 degree turns directed away from the side of the 6-OHDA lesion, reflected hyperstimulation of sensitized postsynaptic dopamine receptors in the denervated striatum. Ipsilateral rotations, defined as turns directed toward the lesion side, were recorded to monitor baseline asymmetry or nonspecific motor activation. Data were expressed as net full turns per time interval, calculated by subtracting ipsilateral turns from contralateral turns, or as total cumulative net rotations per testing session to evaluate the onset, peak effect, and duration of action of the administered compounds and the impact of apelin peptide treatments. We use a pre and post testing paradigm with the virus mediated expression of apelin36 peptides. The hemiparkinsonian mice (WT, D1cre and A_2A_cre) were initially pretested for rotational behavior for 5 days at 25mg/Kg and grouped to receive control GFP, apelin36 or scrambled peptide expressed via AAV infection of the lesioned striata. The mice recovered for 4 weeks post-surgery and tested for L-DOPA (25mg/Kg) induced rotational behavior for another 10 days.

## Statistical analysis

Statistical analyses were performed using StatView (version 5.0.1, SAS Institute Inc.) and GraphPad Prism (version 10, GraphPad Software). Non-parametric assessments were applied to non-normally distributed behavioral rating scales. Specifically, AIM scores were analyzed using the Mann-Whitney U test for non-paired, two-group comparisons and the Wilcoxon matched-pairs signed-rank test for within-subject paired evaluations. Automated rotational behavior was evaluated using a two-way repeated-measures analysis of variance (RM-ANOVA), incorporating treatment (apelin peptide versus vehicle) and testing day as factors, followed by Bonferroni post hoc tests to correct for multiple comparisons across time points. Concentration response curves, kinetic signaling profiles, and baseline normalized luminescence values from real time cAMP accumulation and nanoBiT arrestin recruitment assays were analyzed using GraphPad Prism. Non-linear regression models (four-parameter logistic fits) were used to determine E_max_ and EC_50_ values, and experimental groups were compared using one-way or two-way ANOVA with appropriate post hoc test corrections (such as Bonferroni or Dunnett post hoc analyses) based on the number of comparisons. Quantitative western blot optical density measurements were analyzed using one-way or two-way ANOVA, followed by Bonferroni post hoc tests for pairwise comparisons between groups. Data are presented as mean ± standard error of the mean (SEM), and statistical significance was defined as p<0.05.

## Results

### Dopamine circuitry and L-DOPA induced dyskinesia

The schematic mapping of nigrostriatal circuitry illustrates the progressive transition from a healthy physiological state to Parkinsonian neurodegeneration and the subsequent emergence of dyskinesia following chronic replacement therapy (Figure 1A-C). Under healthy physiological conditions, presynaptic dopaminergic projections originating in the substantia nigra pars compacta supply controlled levels of dopamine to medium spiny neurons in the striatum (Figure 1A). Degeneration of these midbrain dopaminergic neurons in PD leads to profound striatal DA depletion, impairing coordinated motor function. Exogenous replacement therapy with L-DOPA is converted into DA by remaining catecholaminergic terminals, as well as ectopic serotonergic neurons expressing DOPA decarboxylase (Figure 1B). The severe denervation of dopaminergic nerve terminals reduces synaptic dopamine transporter levels, impairing normal presynaptic reuptake mechanisms. Consequently, unregulated dopamine release causes non-physiological synaptic accumulation and non-pulsatile stimulation, which induces hypersensitization of postsynaptic DARs expressed on striatal MSNs (Figure 1B). At the circuit level, striatal output is divided into direct and indirect projection pathways that receive differential inputs from cortical, cholinergic, and raphe nucleus projections (Figure 1C-E). Pro-kinetic direct pathway medium spiny neurons express G_αs/olf_ coupled DA-D_1_R alongside APJR (Figure 1D). Activation of these direct pathway MSNs stimulates intracellular cAMP signaling and PKA cascades, projecting inhibitory GABAergic signals directly to the internal segment of the globus pallidus and substantia nigra pars reticulata to disinhibit thalamocortical execution of movement. Conversely, anti-kinetic indirect pathway MSNs express G_αi/o_ coupled DA-D_2_R, which inhibit AC activity and suppress movement through polysynaptic circuitry involving the external segment of the globus pallidus and subthalamic nucleus (Figure 1E). The pathological sensitization of DARs, predominantly marked by hyperactive D_1_R signaling within direct pathway neurons, creates an aberrant circuit imbalance that drives LID.

**Figure 1:**
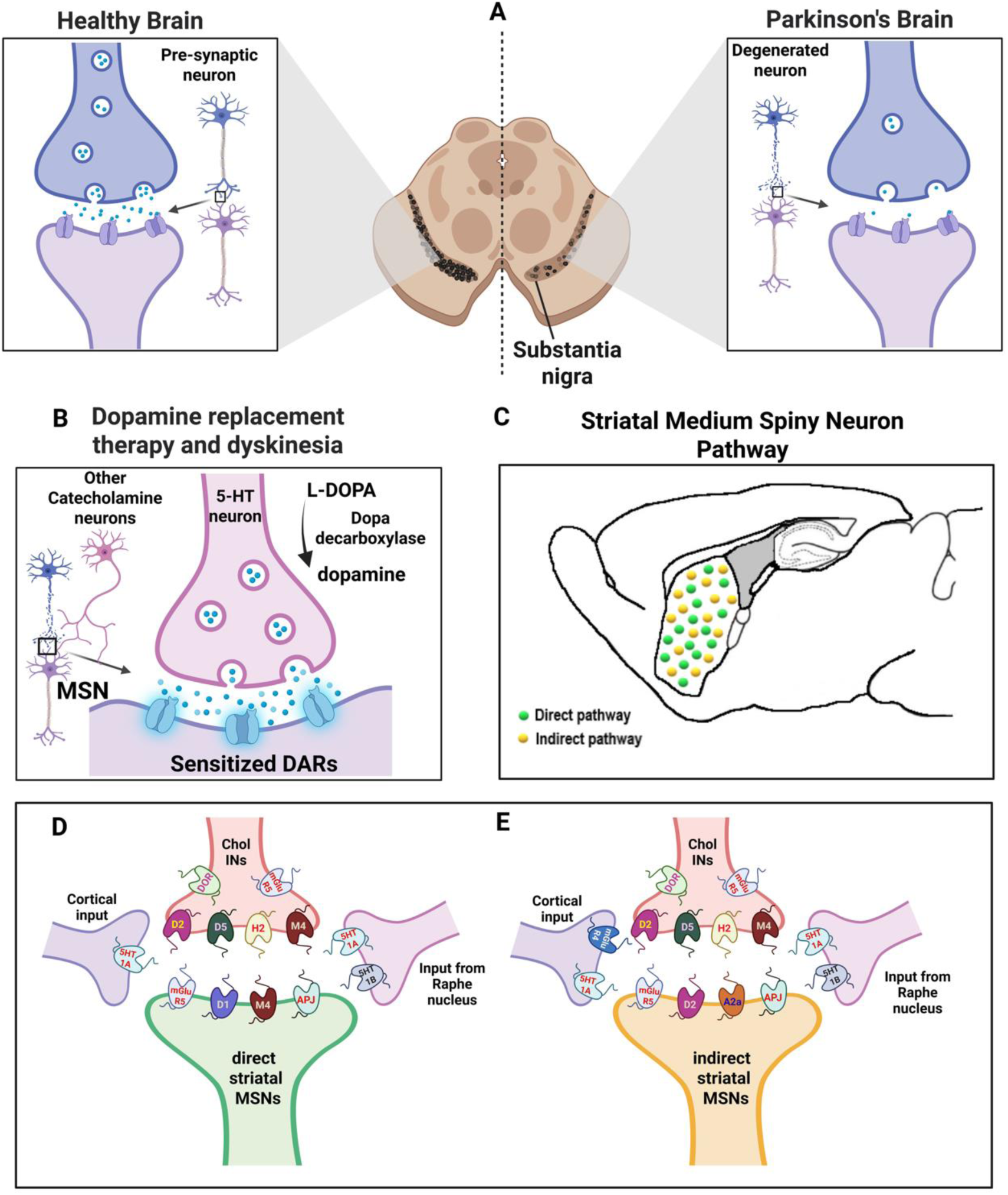
Schematics for Parkinson’s disease (PD) pathology, dopamine replacement therapy with L-3,4-dihydroxyphenylalanine (L-DOPA) and striatal medium spiny neuron (MSN) pathways. **A.** Cartoons depicting healthy and Parkinsonian brain at the level of Substantia nigra where degeneration of dopamine producing presynaptic neurons results in the loss of dopamine supply to striatal MSNs leading to the clinical manifestations of PD. **B.** Modeling of dopamine replacement therapy and L-DOPA induced dyskinesia (LID). L-DOPA is converted in the catecholaminergic neurons to dopamine via the action of the enzyme DOPA decarboxylase. Loss of dopaminergic neurons enables other catecholaminergic neurons such as 5-hydroxytrypamine (5-HT) neurons to aid in the production of dopamine. Dopaminergic denervation also results in the loss of dopamine transporters (DAT) abrogating the reuptake mechanism leading to excess availability of dopamine in the synapses and eventual sensitization of dopamine receptors (DARs) present in the MSNs of the striatal pathway neurons. **C. Dopaminergic projection neurons and circuit balance:** Dopamine acts synergistically across both subpopulations exciting D1-direct MSNs while suppressing indirect MSNs to facilitate coordinated movement. Imbalance between these pathways underlies motor dysfunction in disorders such as Parkinson’s disease (D_1_ hypoactivity / D_2_ hyperactivity) and L-DOPA-induced dyskinesia. **D. Direct Pathway (Pro-kinetic):** Direct pathway medium spiny neurons (direct MSNs) express dopamine D1 receptors (Gα_s/olf_ -coupled) and co-express substance P and dynorphin. Nigrostriatal dopamine activates direct MSNs via D1R, promoting intracellular cAMP production and protein kinase A (PKA) signaling. Direct MSNs project GABAergic inhibitory projections directly to the internal segment of the globus pallidus (GPi) and substantia nigra pars reticulata (SNr). Inhibition of GPi/SNr output disinhibits glutamatergic thalamocortical projections, promoting motor execution and behavioral reinforcement. **E. Indirect Pathway (Anti-kinetic):** Indirect pathway medium spiny neurons (indirect MSNs) express dopamine D2 receptors (G_αi/o_ -coupled) and co-express enkephalin. Nigrostriatal dopamine inhibits indirect MSNs via D2R activation, suppressing adenylyl cyclase activity. Indirect MSNs send GABAergic projections to the external segment of the globus pallidus (GPe), which in turn inhibits the subthalamic nucleus (STN). Glutamatergic excitatory drive from the STN to the GPi/SNr increases basal ganglia inhibitory tone onto the thalamus, suppressing unintended motor programs. Figure 1 was made using Biorender for publication (License agreement number HM2AG4JBKN).

### Dopamine mediated GPCR signaling and its implications on the intracellular pathways

The signaling cross talk and regulatory dynamics between G_αs_ and G_αi/o_ coupled receptors that shape striatal MSN activity was investigated by evaluating intracellular cAMP dynamics using a glosensor real time bioluminescence assay in HEK293 cells. The conceptual framework illustrates that D_1_R activation stimulates G_αs/olf_ proteins to activate AC, converting ATP to cAMP and activating PKA to drive downstream cellular responses (Figure 2A). Conversely, D_2_R stimulation recruits G_αi/o_ proteins to inhibit AC activity, counteracting cAMP generation. In cells expressing DA D_1_R alone, stimulation with 10µM DA produced a robust and sustained peak in cAMP production (Figure 2B). Co-expression of D_2_R along with D_1_R resulted in a significant reduction in cAMP generation over time (p<0.0001 and p<0.05), demonstrating functionally competitive inhibition of AC by G_αi/o_ signaling (Figure 2B). Concentration response evaluations revealed that co-expression of D_2_R shifted the potency of DA, increased approximately eightfold from 0.0029µM for D_1_R alone to 0.0238µM for the D_1_ and D_2_R combination (Figure 2C). The D_1_R and D_2_R expression along with loading control GAPDH is shown in figure 2D. To test whether preactivation or pharmacological blockade of G_αi/o_ coupled receptors alters this balance, cells co-expressing D_1_ and D_2_R were treated with pathway specific ligands prior to DA stimulation (Figure 2E-G). Pre-treatment with the D_2_R agonist quinpirole (10 µM) for 10, 20, or 30 minutes before DA addition progressively disengaged G_αi/o_ mediated suppression, leading to time dependent increases in D_1_R driven cAMP luminescence (p<0.05) (Figure 2E). Similarly, pharmacological inhibition of D_2_R using the antagonist spipirone attenuated G_αi/o_ inhibition, enhancing cAMP accumulation following DA application (p<0.01) (Figure 2G). The expression of D_1_ and D_2_R in the cells for the agonist and antagonist treatments of D_2_R respectively are shown in figures F and H. To determine if this regulatory interaction is selective to dopamine receptors, D_1_R were co-expressed with the muscarinic M_2_R, another G_αi/o_ coupled receptor. Baseline inhibition of D_1_R driven cyclic AMP by muscarinic M_2_R co-expression was markedly weaker than that exerted by DA-D_2_R, though pre-incubation with the agonist carbachol (10µM) for 30 minutes significantly increased subsequent DA stimulated cAMP production (p<0.05) (Figure 2I) and the expression of D_1_R, M_2_R-LgBiT are shown in figure 2J.

**Figure 2:**
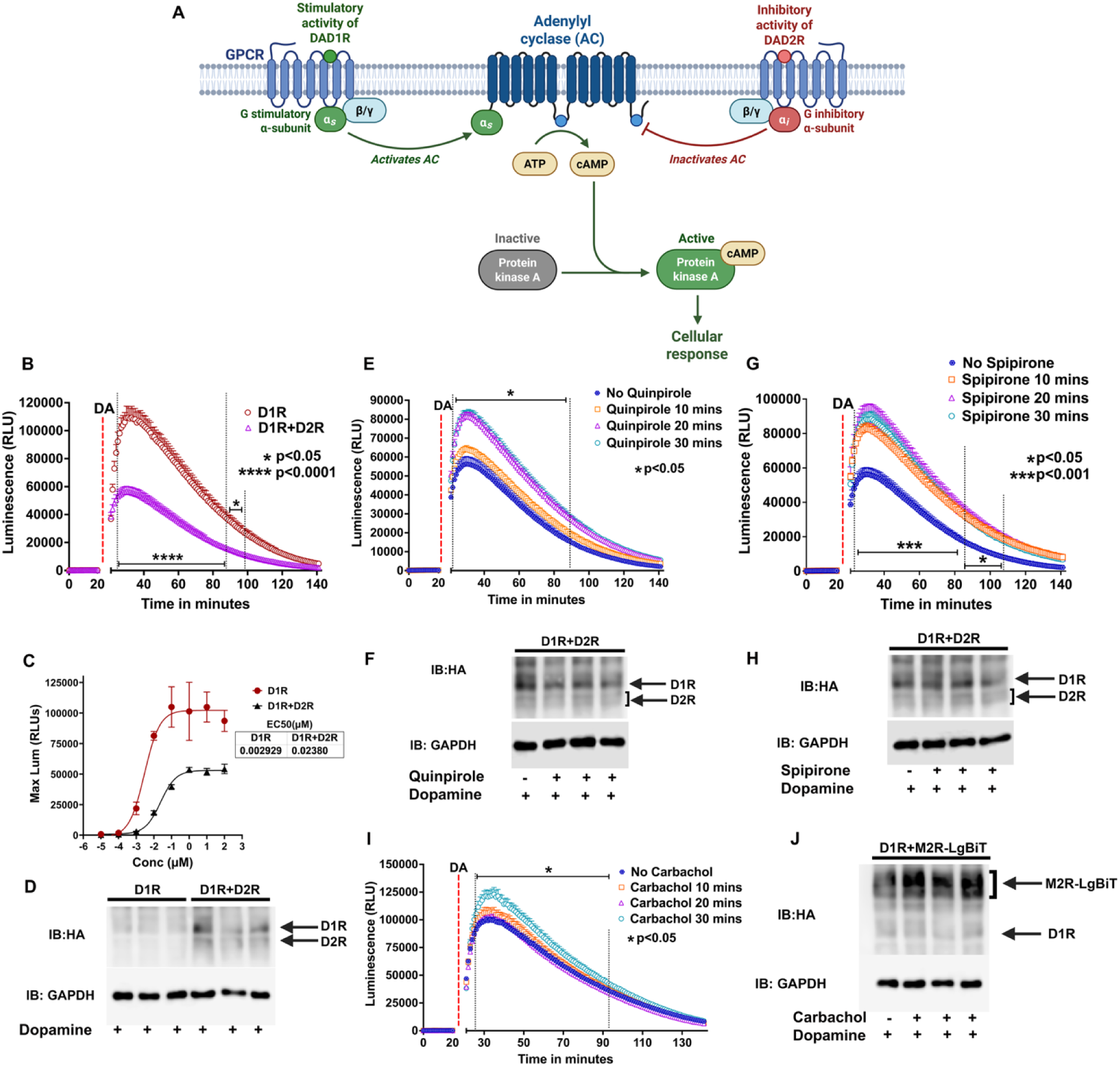
Dopamine mediated GPCR signaling and competitive cAMP assay. **A.** D_1_R activation and PKA/DARPP-32 signaling in dMSNs by dopamine action on DAR-D1 on direct pathway MSNs activate D_1_R-G_αs/olf_ coupling. This stimulates adenylyl cyclase (AC5) and activates intracellular cyclic AMP (cAMP), and subsequent activation of Protein Kinase A (PKA) and the 32-kDa dopamine and cAMP regulated phosphoprotein (DARPP-32) culminating in cellular responses via MAPK and other pathways to drive forward motion. On the other hand, D_2_R (D_2_R-G_αi/o_) activation in indirect MSNs maintains suppression of the PKA/DARPP-32 cascade. **B.** cAMP assay using Glosensor22F with expression of either dopamine D_1_R or D_1_R + D_2_R in HEK293 cells. Basal activity measured for 20minutes just before the addition of the ligand for stimulation. Stimulation using dopamine at a concentration of 10µM of the D1R produces robust cAMP activation which is abrogated or significantly reduced in the presence of D_2_R which also competes for the ligand inhibiting adenylate cyclase activation via G_αi_ reducing cAMP production. The cAMP activation was used as a competitive assay to differentiate the involvement of GPCRs other than DARs in the direct and indirect neuronal pathways. Plots indicate the relative luminescence over time showing the increase in the cAMP production and were from 8 individual wells of the 96 well format averaged and presented as mean ± S.E.M. The background luminescence was read 20 minutes before the addition of the ligand. Statistical analysis using two-way ANOVA showed significant reduction (p<0.0001 and p<0.05 over time respectively as indicated) in the cAMP production in the cells expressing both D_1_ and D_2_R than D_1_R expression alone. **C.** The graph shows the EC_50_ of dopamine for D1R alone and D1R+D2R expression in HEK293 cells using escalating dose of dopamine for stimulation in a glosensor cAMP assay. The max luminescence for each dose range averaged from 4 separate well readings were used for plotting and the EC_50_ was calculated from the slope of the curve. The EC_50_ was reduced 8-fold, 0.0029 vs 0.023 for D_1_R alone compared with D_1_R+D_2_R respectively. **D** shows the expression of HA tagged D1R and D2R in the cAMP assay with dopamine stimulation. GAPDH is used as a loading control. **E.** Pre-activation using D_2_R agonist quinpirole in a time dependent manner results in increased cAMP activation via the D_1_R when D_1_R and D_2_R are co-expressed in HEK293 cells. The activation of cAMP via stimulation of D_1_R using dopamine was used as a readout for this competitive assay between the DARs. Activation of D_2_R using the agonist quinpirole (10µM) before stimulation with dopamine disengages the D_2_R and promotes cAMP production via the D_1_R. **F** shows the expression of HA tagged D_1_ and D_2_R with and without D_2_R agonist quinpirole respectively and dopamine agonists. GAPDH is used as a loading control. **G.** Antagonism of D_2_R with spipirone (10µM) completely disengages the D_2_R when co-expressed with D_1_R in the cAMP activation assay. Dopamine alone suppresses the cAMP activity when D_1_R and D_2_R are co-expressed due to the competitive inhibition by D_2_R, however when the activity of D_2_R is blocked by the antagonist cAMP production is promoted via D1R upon stimulation by dopamine. **H** shows the expression of HA tagged D_1_ and D_2_R with and without D_2_R antagonist spipirone respectively and dopamine agonists. GAPDH is used as a loading control. **I.** The competitive cAMP assay was performed with another G_αi_ coupled GPCR, the muscarinic 2 receptor (M_2_R) co-expressed along with D_1_R in HEK293 cells. M_2_R co-expression with D_1_R had a much less effect on the inhibition of cAMP activation mediated by D_1_R, however pre-incubation with carbachol (10µM) for 30 minutes before the addition of dopamine did increase the cAMP activation significantly than no carbachol and carbachol addition for 10 and 20 minute time points. Taken together the data suggests the opposing effects of G_αs_ and G_αi_ coupled GPCRs on adenylate cyclase expressed together and their effects on cAMP activation and cellular signaling responses. **J.** Western blot expression for dopamine D_1_R, muscarinic M_2_R-LgBiT tagged with HA and GAPDH in HEK293 cell transfection experiments. Figure 2A was made using Biorender for publication (License agreement number ES2AG4K6BD).

### Homologous desensitization – a mechanistic pathway in LID

The mechanisms governing receptor desensitization and arrestin recruitment, were evaluated by real time kinetics and spatial localization of DA-D_1_R interactions with arrestin isoforms (Figure 3). Agonist binding initiates homologous desensitization via G protein coupled receptor kinase (GRK) phosphorylation, driving arrestin translocation to the plasma membrane to terminate canonical G protein signaling and trigger receptor endocytosis, degradation, or recycling (Figure 3A). Using the nanoBiT split luciferase complementation assay, we monitored the interaction between D_1_R-LgBiT and either arrestin2-SmBiT or arrestin3-SmBiT in HEK293 arrestin2/3dko cells (Figure 3B-D). Addition of 10µM DA to cells expressing D_1_R-LgBiT and arrestin2-SmBiT induced a rapid and robust increase in bioluminescence within minutes, indicating immediate recruitment of arrestin2 to the active receptor, followed by a gradual decay over 60 minutes (Figure 3C). A similar dynamic profile was observed in cells expressing D_1_R-LgBiT and arrestin3-SmBiT, where DA stimulation elicited a marked increase in luminescent signal, reaching high recruitment levels that steadily declined over time (Figure 3D). Confocal laser scanning microscopy validated the spatial translocation of arrestin isoforms to the plasma membrane in dko cells (Figure 3E-H). Under basal non stimulated conditions, GFP tagged arrestin2 (Figure 3E) and arrestin3 (Figure 3G) exhibited diffuse cytosolic localization, whereas HA tagged D_1_R localized predominantly to the plasma membrane. Stimulation with 10µM DA for 15 minutes induced prominent translocation of both arrestin2 GFP (Figure 3F) and arrestin3 GFP (Figure 3H) to the plasma membrane, displaying strong co-localization with membrane bound DA-D_1_R.

**Figure 3:**
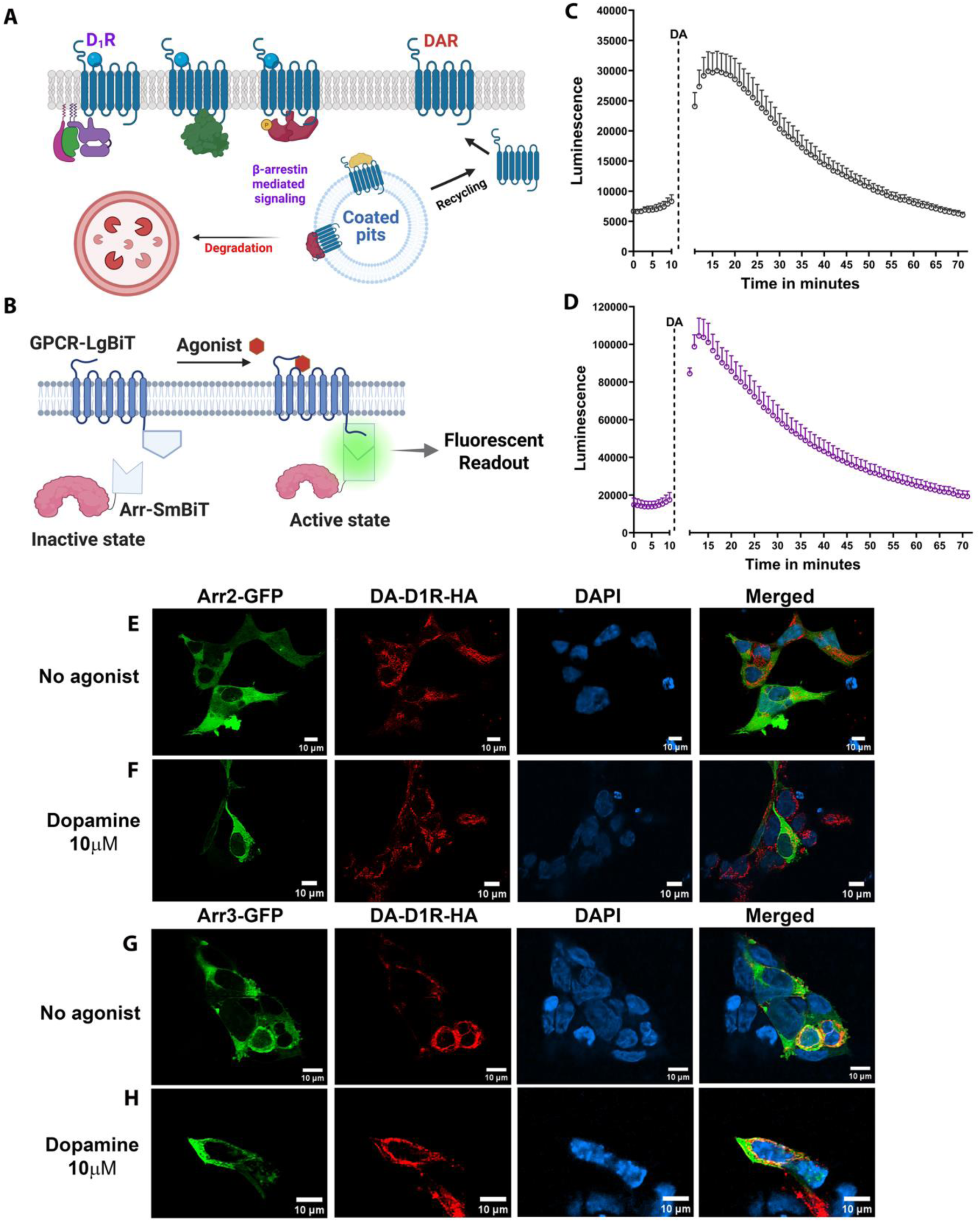
**A.** Classical Homologous Desensitization: Agonist binding to a G protein-coupled receptor (GPCR) triggers a conformational change that promotes G protein coupling and downstream intracellular signaling. To prevent sustained overstimulation, active receptors undergo homologous desensitization initiated by G protein-coupled receptor kinases (GRKs). GRKs selectively phosphorylate serine and threonine residues on the intracellular loops and carboxyl-terminal tail of the agonist-bound receptor. Phosphorylation creates a high-affinity binding site for β-arrestin (β-arrestin-1/2), which translocates from the cytosol to the plasma membrane. β-Arrestin sterically hinders further G protein coupling (uncoupling), terminating canonical G protein signaling. Subsequently, membrane-bound β-arrestin links the receptor to clathrin-coated pits via interactions with clathrin and AP-2, targeting the receptor for endocytosis, degradation, or recycling. **B.** NanoBiT bioluminescence complementation assay for arrestin recruitment: Real-time monitoring of β-arrestin recruitment to the plasma membrane GPCR is quantified using the NanoLuc Binary Technology (NanoBiT) split-luciferase system. The large subunit of NanoLuc luciferase (LgBiT, ∼18 kDa) is genetically fused to the C-terminus of the target GPCR, while the complementary small peptide subunit (SmBiT, 11 amino acids, low affinity) is fused to the N or C terminus of β-arrestin. Under basal conditions, LgBiT and SmBiT remain dissociated due to their low intrinsic affinity, generating negligible background luminescence. Upon agonist stimulation, GRK mediated receptor phosphorylation drives the rapid translocation of SmBiT β-arrestin to the LgBiT -GPCR at the plasma membrane. Proximity-induced structural complementation of LgBiT and SmBiT reconstitutes functional NanoLuc enzyme activity. In the presence of the Coelenterazine H (Coe H) substrate, this interaction yields a bright, real-time luminescent signal proportional to the dynamics and kinetics of β-arrestin recruitment. **C.** Graphical presentation of the signaling response for the nanoBiT assay with D_1_R-LgBiT and arrestin2-SmBiT in HEK293 arrestin2/3 double knockout (arr2/3-dko) cell lines. The luminescence intensity was plotted over time with basal activity read for 10 minutes after the addition of substrate Coe H followed by the addition of agonist dopamine (10µM) to activate D_1_R for another 1 hour. Addition of the ligand induces a robust recruitment of arrestin2-SmBiT to the membrane producing an intense luminescence signal as a result of GPCR-arrestin interaction. **D.** Graphical presentation for the signal output of nanoBiT assay with D_1_R-LgBiT and arrestin3-SmBiT in HEK293 arr2/3-dko cell line. The experiment was conducted under similar conditions to arrestin2 nanoBiT assay with readings averaging from 4 wells in both the experiments and repeated thrice for statistical analysis. **E.** Confocal images for arrestin2-GFP and D_1_ receptor in HEK293 arr2/3-dko cells (green channel for GFP, red channel for HA tagged to D1R, blue channel for nuclear stain DAPI and merged composite images show colocalization of the signal). The arrestin2-GFP expression was found diffused throughout the cytosol under basal conditions and the receptor expression was more localized to the membrane. Note: Native fluorescence was used for GFP signal and the HA signal was amplified using HA primary antibody which was subsequently enhanced using alexafluor 568 secondary antibody after fixation of the cells using paraformaldehyde. **F.** Confocal images for arrestin2-GFP recruitment to the D_1_ receptor in HEK293 arr2/3-dko cells upon stimulation with agonist dopamine (10µM). Agonist stimulation results in the translocation of arrestin2 to the receptor with intense signal observed at the membrane. **G.** Confocal images for arrestin3-GFP and D_1_ receptor in HEK293 arr2/3-dko cells (green channel for GFP, red channel for HA tagged to D1R, blue channel for nuclear stain DAPI and merged composite images show colocalization of the signal). Similar to arrestin2, arrestin3-GFP expression was found diffused throughout the cytosol under basal conditions and the receptor was localized to the membrane. **H.** Confocal images for arrestin3-GFP recruitment to the D_1_ receptor in HEK293 arr2/3-dko cells upon stimulation with agonist dopamine (10µM). Agonist stimulation results in the translocation of arrestin3 to the receptor with intense signal colocalization observed at the membrane. Scale bar in the confocal images denotes 10µm. Figure 3A and B was made using Biorender for publication (License agreement number NI2AG4JWOO).

### Apelin receptor synergistically competes with DAR in dampening down signaling

APJ receptor cross-regulation of D_1_R intracellular signaling was determined by competitive cAMP assays, nanoBiT arrestin recruitment kinetics, and confocal imaging in HEK293 cell models (Figure 4). Coexpression of G_αi/o_ coupled APJR with D_1_R significantly attenuated 10µM DA stimulated cAMP accumulation over time (p<0.0001) following pre-treatment with 10µM apelin13 (Figure 4A). Concentration response analyses revealed that apelin13 treatment shifted the potency of DA, increasing the EC_50_ nearly four-fold from 0.0452µM in non-treated controls to 0.1773µM in apelin treated cells (Figure 4B). Evaluation of arrestin recruitment using nanoBiT assays in dko cells demonstrated that 10µM apelin13 induced robust, immediate recruitment of both arrestin2-SmBiT and arrestin3-SmBiT to APJR-LgBiT (p<0.01 and p=0.0001 relative to PBS control), with arrestin2 showing approximately two-fold higher peak recruitment fold change compared to arrestin3 (Figure 4C, D). In triple cotransfection experiments (APJR-DUET, D_1_R-LgBiT, and arrestin-SmBiT), pre-incubating cells with apelin13 for 10, 20, or 30 minutes prior to DA application completely abrogated DA induced recruitment of arrestin2 (p<0.0001) and significantly blunted arrestin3 recruitment to the D_1_R (p<0.0001) (Figure 4E, F). Confocal laser scanning microscopy shows diffuse cytosolic distribution of GFP-tagged arrestin2 (Figure 4G) and arrestin3 (Figure 4I) under basal conditions. Apelin stimulation results in translocation of arrestins to APJR at the plasma membrane (Figure 4H, J). We further evaluated the longer isoform, apelin36, using vector mediated peptide expression (pAAV-EF1α-apelin36-IRES-GFP) (Figure 4K-N). Recombinant apelin36 expression markedly inhibited DA induced cAMP generation compared to a scrambled apelin36 control (p<0.0001 and p<0.05) (Figure 4K). This expression shifted the dopamine concentration-response curve, increasing the DA EC_50_ approximately three-fold from 0.0068µM (scrambled) to 0.0194 µM (wild type apelin36) (Figure 4L). Apelin treatment reduced D1R-mediated cAMP responses and altered D1R-arrestin recruitment in HEK293 cells, whether this reflects competition for arrestins, changes in receptor trafficking, or other forms of signaling crosstalk remains unclear. The receptor expressions of APJR, D_1_R, APJR-LgBiT, D_1_R-LgBiT and arr2-SmBiT and arr3-SmBiT for the cAMP and nano-BiT respectively are shown in figures 4O, P and Q.

**Figure 4:**
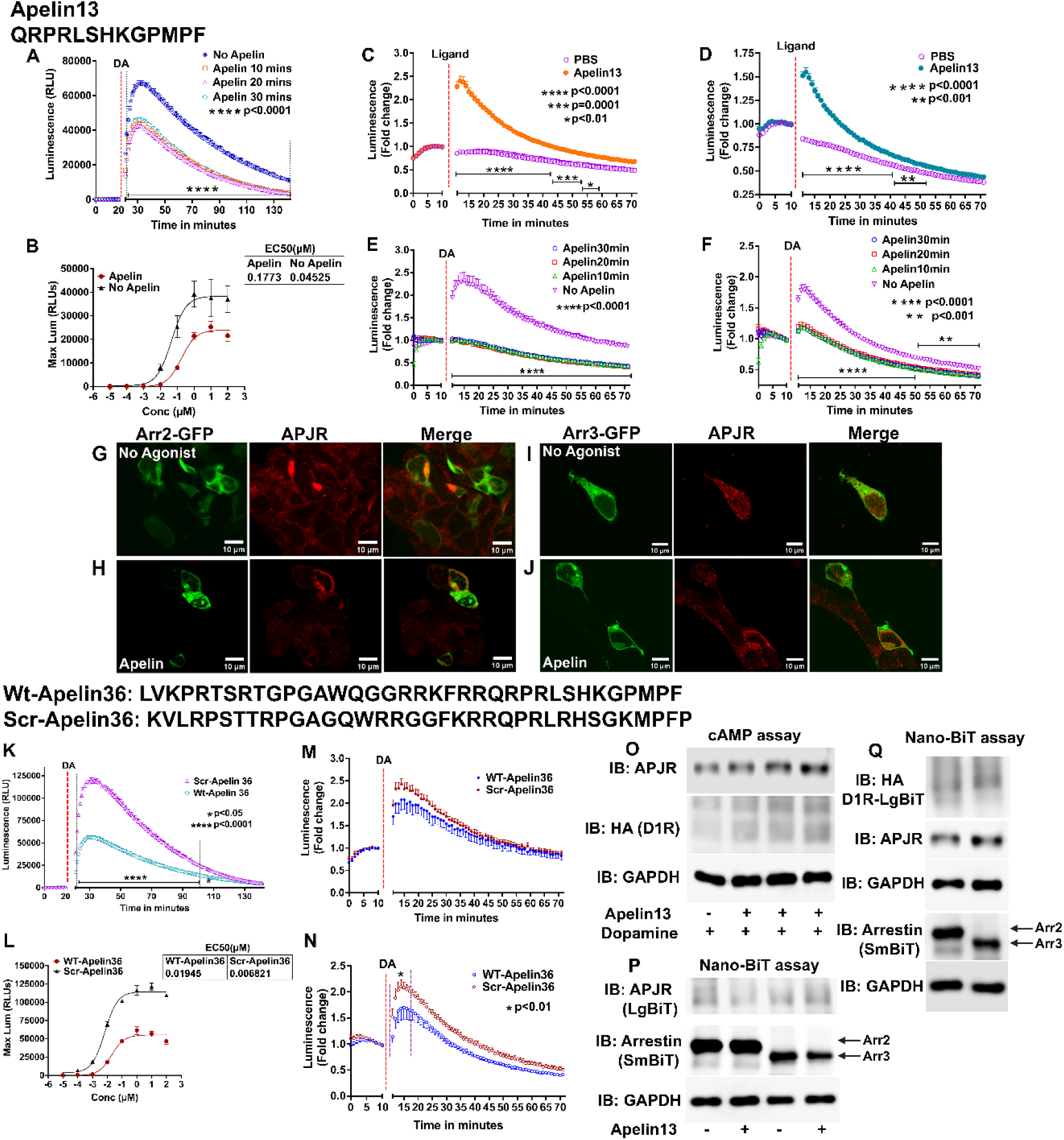
Signaling responses with apelin peptides in HEK293 cells. **A.** Competitive cAMP assay using Glosensor22F with co-expression of DA-D_1_R and APJR in HEK293 cells. Stimulation using DA at a concentration of 10µM of the D1R produces robust cAMP activation which is significantly reduced (p<0.0001, one-way ANOVA) in the presence of APJR pretreated with apelin13 peptide ligand (10µM). cAMP assay plots indicate the relative luminescence presented as mean ± S.E.M. APJR is a G_αi_ coupled GPCR which inhibits AC activation via G_αi_ reducing cAMP production. The co-expression of APJR along with D_1_R in the cells strongly abrogates the cAMP responses mediated via the D1R blunting the signaling mediated via PKA. **B.** The graph shows the EC_50_ of dopamine for D_1_R+APJR expression in HEK293 cells using escalating doses of DA with and without apelin13 for stimulation in a glosensor cAMP assay. The max luminescence for each dose range averaged from 4 separate well readings were used for plotting and the EC_50_ was calculated from the slope of the curve with the experiments replicated thrice. The EC_50_ increased about 4-fold, 0.177 vs 0.045 µM for apelin compared with no apelin respectively. This relates to the suppression of D_1_R signaling responses in the presence of APJR due to competitive inhibition of the AC activation by APJR thereby reducing the overall responses due to D_1_R. **C.** Graphical presentation of the signaling response for the nanoBiT assay with APJR-LgBiT and arrestin2-SmBiT in HEK293 arr2/3-dko cell lines. The luminescence intensity was plotted over time with basal activity read for 10 minutes after the addition of substrate Coe H followed by the addition of apelin13 (10µM) to activate APJR for another 1 hour. Addition of the ligand induces a robust recruitment of arr2-SmBiT to the membrane producing an intense luminescence signal as a result of GPCR-arrestin interaction and is presented as fold change to the basal activity which was significant over time (p<0.01, p=0.0001 and p<0.0001) using one-way ANOVA. Phosphate buffered saline (PBS) used as an agonist control did not elicit any arrestin2 recruitment. **D.** Graphical presentation of the signaling response for the nanoBiT assay with APJR-LgBiT and arr3-SmBiT in dko cell lines with apelin13 (10µM). Apelin13 induces the recruitment of both arr2 and arr3 to the receptor, though the recruitment of arr2 was twofold more than that of arr3. The statistical analysis was performed by one-way ANOVA for luminescence fold change from 4 well readings with the experiments repeated thrice and averaged for graphical presentation in C and D. p values shows the statistical difference between the peptide agonist and PBS control over time. **E.** Graphical presentation of the signaling response for the nanoBiT assay with coexpression of APJR-DuET, D_1_R-LgBiT and arr2-SmBiT in dko cell lines. The cells were pretreated with apelin13 (10µM) for 10, 20 and 30 minutes to target APJR before the addition of DA. Coe H was added 10 minutes before the stimulation of D1R with DA (10µM) to all the wells. The wells without apelin pretreatment but with dopamine served as positive control. The luminescence intensity was plotted over time for all the experimental groups. Addition of DA induced a robust recruitment of arr2-SmBiT to the membrane producing an intense luminescence signal only in the wells not pretreated with apelin. Apelin pretreatment before the addition of DA abrogated the recruitment of arr2-SmBiT to the D_1_R strongly engaging arr2 as a result of competitive interaction. p values show statistical difference between with and without apelin pretreatment over time. **F.** Graphical presentation of the signaling response for the nanoBiT assay with coexpression of APJR-DuET, D_1_R-LgBiT and arr3-SmBiT in dko cell lines. The cells were treated in a similar way as with arr2-SmBiT experiment. Apelin pretreatment before the addition of DA diminished the recruitment of arr3-SmBiT to the D_1_R similar to arr2 as a result of competitive interaction. p values show statistical difference between with and without apelin pretreatment over time. **G.** Confocal images of basal expression of arr2-GFP and APJR in dko cells (green channel for GFP, red channel for APJR and merged composite images show colocalization of the signal). **H.** Confocal images of apelin13 treatment and recruitment of arr2-GFP to APJR in dko cells (green channel for GFP, red channel for APJR and merged composite images show colocalization of the signal). **I.** Confocal images of basal expression of arr3-GFP and APJR in dko cells (green channel for GFP, red channel for APJR and merged composite images show colocalization of the signal). **J.** Confocal images of apelin13 treatment and recruitment of arr3-GFP to APJR in dko cells (green channel for GFP, red channel for APJR and merged composite images show colocalization of the signal). **K.** Competitive cAMP assay with co-expression of DA-D_1_R, APJR and pAAV-EF1α-GFP-IRES-apelin36 or scrambled apelin36 peptide constructs in wild type HEK293 cells. The experimental paradigm used the cell expression of the apelin36 peptides and DA (10µM) was used to stimulate the D_1_R. Apelin36 expression in HEK cells strongly inhibited the cAMP induced by DA stimulation of D_1_R than the expression of scrambled apelin36 peptide under similar DA stimulation of the D_1_R. The reduction of the cAMP could be attributed to the stimulation of APJR which is coupled to G_αi_ which inhibits AC activity upon stimulation. cAMP assay plots indicate the significantly reduced (p<0.0001 and p<0.05, one-way ANOVA) luminescence over time in apelin36 expressing cells than scrambled apelin36 and the values were presented as mean ± S.E.M. **L.** The graph shows the EC_50_ of DA for D_1_R+APJR expression in HEK293 cells using escalating doses of DA with endogenous expression of apelin36 or scrambled apelin36 peptide in a glosensor cAMP assay. The EC_50_ increased about 3 fold, 0.0194 vs 0.00682µM for apelin36 compared with scrambled apelin36 expression respectively. This relates to the suppression of D_1_R signaling responses in the presence of APJR due to competitive inhibition of the AC activation by APJR thereby reducing the overall responses due to D_1_R. **M.** Graphical presentation of the signaling response for the nanoBiT assay with coexpression of APJR-DuET, D_1_R-LgBiT, arr2-SmBiT and either pAAV-apelin36 or scrambled apelin36 constructs in dko cells stimulated with DA (10µM). There was no difference between apelin and scrambled peptide expression in the recruitment of arrestin2 when DA was used to stimulate D_1_R. **N.** Graphical presentation of the signaling response for the nanoBiT assay with coexpression of APJR-DuET, D_1_R-LgBiT, arr3-SmBiT and either pAAV-apelin36 or scrambled peptide constructs in dko cells stimulated with DA (10µM). Expression of apelin36 peptide targeting APJR rather than the scrambled peptide was able to significantly inhibit arrestin3 recruitment when D_1_R was targeted using dopamine. **O.** Western blot pictures for APJR and HA tagged D_1_R in cAMP assay, GAPDH was used as a loading control. **P.** Western blot pictures for APJR, arrestin2-SmBiT, arrestin3-SmBiT and GAPDH in a nano-BiT assay and **Q.** western blot pictures for the expression of HA tagged D1R, APJR and arrestin (2 and 3)-SmBiT in a nano-BiT assay. GAPDH is used as a loading control for the blots.

### In vivo expression of apelin peptide ameliorates LID

The therapeutic efficacy of *in vivo* apelin peptide expression in ameliorating LID was investigated utilizing a hemiparkinsonian mice model established via unilateral 6-OHDA lesioning of the MFB (Figure 5). Stereotaxic infusion of 6-OHDA selectively depleted dopaminergic projections, resulting in severe loss of tyrosine hydroxylase (TH) immunoreactivity within the ipsilateral striatum and substantia nigra pars compacta relative to the intact contralateral hemisphere (Figure 5A, B). To assess behavioral sensitization and motor responses, hemiparkinsonian wildtype mice were treated with daily administration of L-DOPA (25 mg/kg) across a 5 day pretesting baseline period (Figure 5C). Mice subsequently received adeno-associated virus injections targeting the ipsilateral striatum to express wildtype apelin36, scrambled apelin36, or control GFP (Figure 5D). Following a 4 week surgical recovery period, chronic L-DOPA administration was reinitiated for 10 consecutive days of post-AAV testing (Figure 5C). Control mice expressing GFP or scrambled apelin36 displayed progressive behavioral sensitization, manifesting as escalating contralateral rotational behavior (p<0.0001) (Figure 5C). In contrast, mice expressing recombinant apelin36 exhibited significant suppression of L-DOPA-induced motor sensitization, maintaining low contralateral rotations throughout the post-testing regimen (p<0.0001) (Figure 5C). The AAV mediated co-cistronic expression of GFP along with TH expression with GAPDH protein control is shown in figure 5D. Immunohistochemical analysis confirmed widespread transduced expression and cell type localization within the lesioned striatum (Figure 5E-O). Low power panoramic cross sections demonstrated selective GFP in the lesioned striatum (Figure 5E), which colocalized extensively with FoxP, a marker for striatal MSNs (Figure 5F, G). High magnification confocal imaging verified that co-cistronic GFP signal colocalized with FoxP positive MSNs in both apelin36 (Figure 5H, J and L) and scrambled apelin36 treated groups (Figure 5I, K and M). Further high power confocal evaluation demonstrated robust intracellular co-expression of GFP alongside FLAG-tagged apelin36 (Figure 5N) or FLAG-tagged scrambled apelin36 (Figure 5O) within lesioned medium spiny neurons.

**Figure 5:**
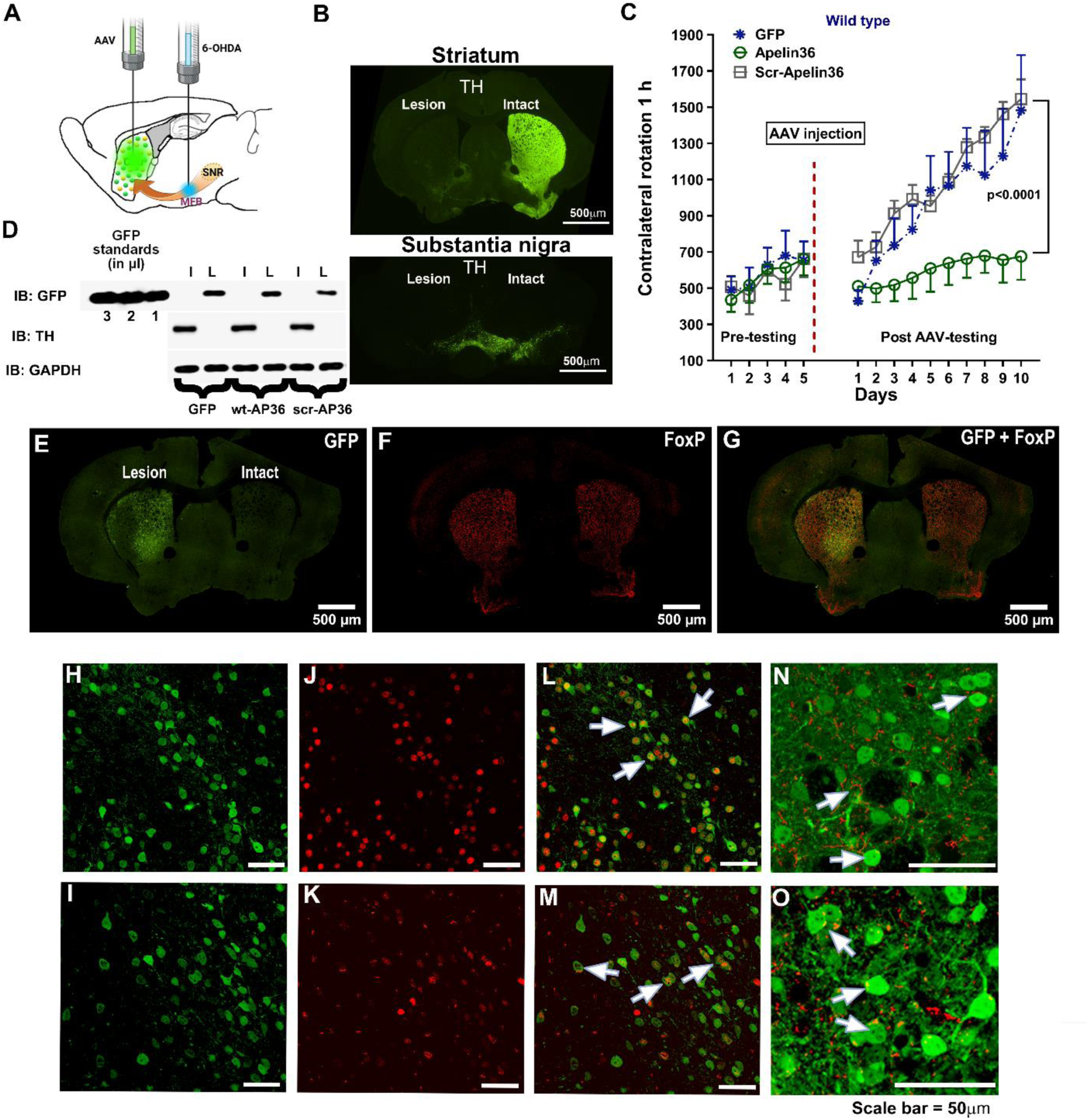
**A.** Schematic of experimental set up in mouse showing unilateral lesion using 6-hydroxydopamine into median forebrain bundle and AAV injection into the striatum of the ipsilateral side of the brain. **B.** Immunohistochemistry for TH expression at the level of striatum and substantia nigra in hemiparkinsonian mouse model. **C.** Contralateral rotational studies in hemiparkinsonian mouse model with AAV mediated expression of apelin36 and scrambled apelin36 peptide with AAV-GFP injected mice serving as positive control. Wildtype hemiparkinsonian mice were chronically treated with L-DOPA (25mg/Kg) for 5 days to measure rotational behavior and were grouped for AAV injection into the ipsilateral side of the lesion and post tested after 4 weeks of surgical recovery with L-DOPA for another 10 days. The increase in contralateral rotational behavior for the experimental groups is shown. To evaluate the effect of viral vector-mediated peptide expression on rotational asymmetry, contralateral rotations (1h) were assessed across post-AAV testing days in wild-type mice treated with GFP, Apelin36, or Scr-Apelin36. A two-way repeated measures ANOVA revealed a significant main effect of treatment (*F*_2,15_ = 5.47, *p*=0.0164), a significant main effect of Time/Days (*F*_9,135_ = 20.40, p<0.0001), and a significant Treatment × Time interaction (*F*_18,135_ = 2.88, *p*=0.0003). Post-hoc analyses demonstrated that apelin36 treatment significantly ameliorated contralateral rotational behavior compared to both GFP and scr-apelin36 control groups over the 10-day post-testing period (*p<*0.0001). **D.** shows the western blot for GFP and TH expression and GAPDH in the hemiparkinsonian model mice (6-OHDA lesion). **E, F** and **G** shows the panoramic view of mouse brain coronal section at the level of striatum for the AAV mediated expression of GFP (E) in the lesioned side, co-stained with FoxP to label dopaminergic medium spiny neurons (F) and composite image of GFP in FoxP expressing neurons (G). **H-I.** High magnification of the lesioned striata showing the AAV mediated expression of GFP (co-cistronic) along with apelin36 peptide and scrambled apelin36 respectively, **J-K.** FoxP expression in the striata and **L-M.** Composite image of GFP and FoxP in the apelin36 and scrambled apelin36 injected mice striata respectively. White arrows indicate co-localization. **N-O.** High magnification confocal images showing co-expression of GFP (green) and Flag tagged apelin36 (N) and scrambled apelin36 (O) respectively in the lesioned striatum. White arrows indicate co-localization. Scale bar equals 50µm in images H through O. Figure 5A was made using Biorender for publication (License agreement number OB2AGE3SSG).

### Translational approaches using apelin peptide via intranasal administration

The translational potential of apelin peptide delivery in mitigating LID was evaluated using behavioral testing in hemiparkinsonian mice. Mice underwent 4 AIM testing sessions across 12 days, followed by a 4 week drug washout period and a 10 day L-DOPA rotation protocol (Figure 6A). Unilateral lesion using 6-OHDA (Figure 6B) was followed with a 5 week post-surgery recovery period. Liposomal apelin13 formulations were administered via non-invasive intranasal delivery 30 minutes prior to twice daily L-DOPA administration, while control animals received empty liposomes. Intranasal liposomal apelin13 administration significantly attenuated the severity of AIMs across all evaluated testing sessions compared to empty liposome controls (p<0.012) (Figure 6C). In contralateral rotational tests following chronic 10 day L-DOPA challenge, intranasal apelin13 pre-treatment significantly suppressed total net contralateral rotations measured over 1 hour compared to control mice receiving empty liposomes (p=0.0051 RM-ANOVA) (Figure 6D). Sub-score evaluation demonstrated that apelin13 treatment markedly reduced total axial, limb, and orolingual abnormal involuntary movement scores across sessions 1 through 4 (p<0.01) (Figure 6E). Analysis of axial, limb, and orolingual movement dynamics in 20 minute time bins revealed a sustained reduction in dyskinetic behaviors during session 1 (Figure 6F) and session 4 (Figure 6G) in apelin13 treated mice compared to control animals. Evaluation of 5 minute time bins for rotations demonstrated progressive motor sensitization in control animals from Day 1 (Figure 6H) through Day 5 (Figure 6I) and Day 9 (Figure 6J). Apelin13 treatment prevented this time dependent sensitization, maintaining stable low rotational counts that differed significantly from control values on Day 5 (p<0.0001) and Day 9 (p=0.0019) (Figure 6I, J).

**Figure 6:**
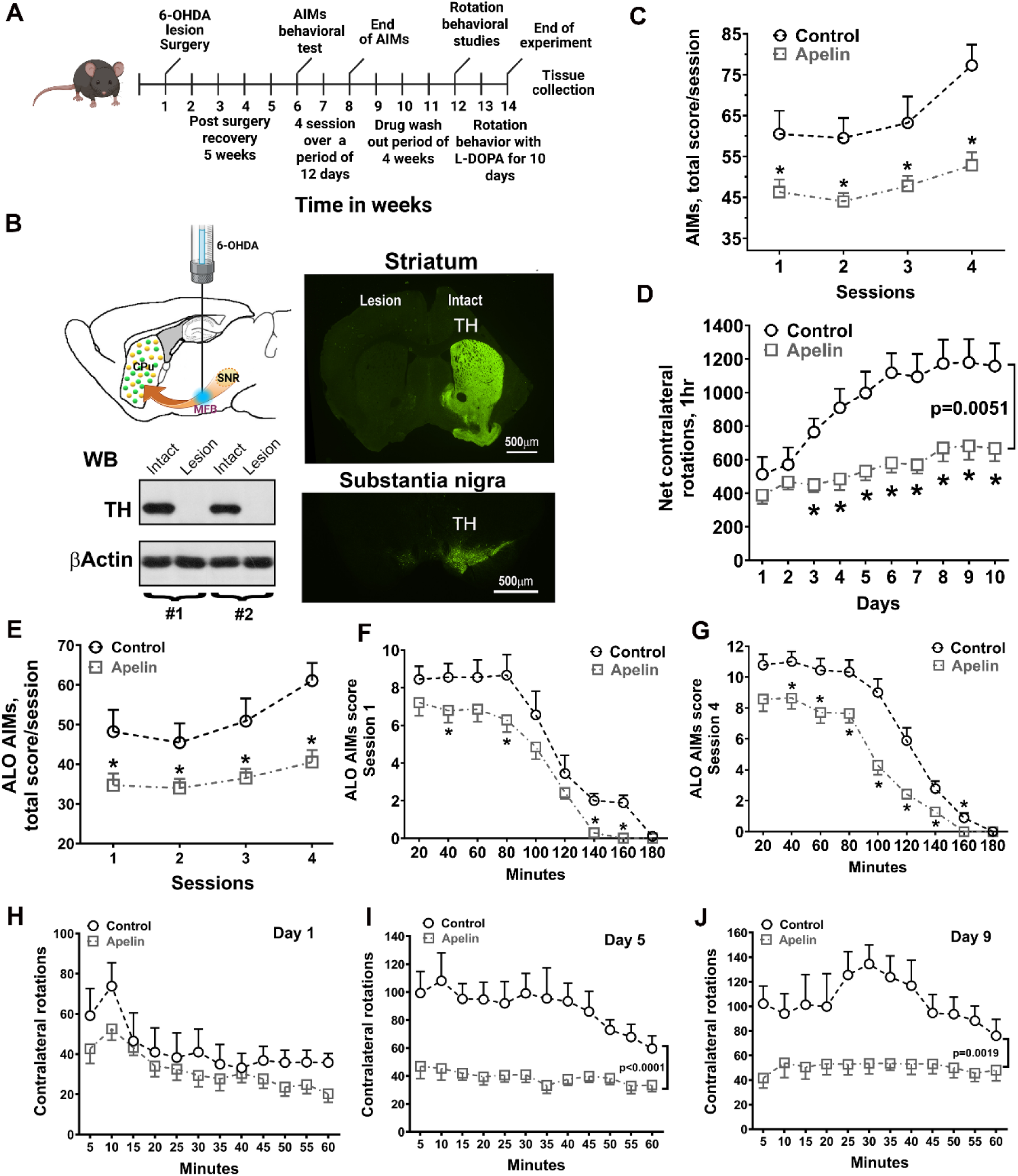
Translational studies of intranasal administration of apelin13 peptide in hemiparkinsonian mouse model. **A.** Timeline for translational behavioral study in mice. **B.** Mouse brain sagittal section showing the placement of the syringe at the level of median forebrain bundle (MFB) for the infusion of 6-OHDA. Pathway arrow shows the depiction of dopaminergic nerve fiber bundle from substantia nigra (SNR) to caudate putamen (striatum). The TH expression after 6-OHDA infusion is shown by western blot and immunohistochemistry at the level of striatum and substantia nigra respectively. **C.** Graph showing total abnormal involuntary movements (AIMs) scores per session between control and apelin groups. The control group consists of intranasal administration of liposomes without peptide. The therapeutic efficacy of apelin-13 administration on dyskinesia severity, AIMs scores were compared between apelin13-infused (n=14) and control (n=10) subjects across Sessions 1 through 4 using Mann-Whitney *U* tests. Apelin13 administration significantly attenuated total AIMs scores compared to the control group across all evaluated sessions. Specifically, Mann-Whitney analyses revealed significant reductions in AIMs scores for Session 1 (U=31.50, Z = -2.25, p=0.0242), Session 2 (U=29.50, Z= -2.37, p=0.0177), Session 3 (U=35.50, Z= -2.02, p=0.0434), and Session 4 (U=12.00, Z= -3.40, p=0.0007). **D.** Graph showing contralateral rotational behavior after chronic L-DOPA administration for 10 days in control and apelin peptide treatment groups. The apelin peptide was delivered via intranasal administration 30 minutes before the administration of L-DOPA twice daily. The rotational behavior was recorded after the daily morning administration after L-DOPA. The control mice received blank liposomal formulations without the peptide. The contralateral rotational behavior decreased significantly in the apelin peptide treatment group than the control mice receiving blank liposomal formulations. the therapeutic impact of Apelin infusion on rotational behavior over an extended testing period, contralateral rotations were evaluated across 10 post-treatment days in Apelin-infused (n=11) and control (n=11) mice. A two-way repeated measures ANOVA revealed a significant main effect of Treatment (F_1,20_ = 9.87, p=0.0051), a significant main effect of Days (F_9,180_ = 16.59, p<0.0001), and a significant Treatment × Days interaction (F_9,180_ = 5.95, p<0.0001). Post-hoc evaluation demonstrated that Apelin-infused subjects exhibited significantly lower rotational asymmetry compared to the control group across the testing timeline, particularly diverging as control rotations increased over consecutive days. **E.** Graph showing total axial, limb and orolingual (ALO-AIMs) scores per session between control and apelin groups. To assess the therapeutic efficacy of apelin on dyskinesia severity, ALO AIMs scores were compared between apelin-infused mice (n=14) and control subjects (n=10) across testing sessions using Mann-Whitney *U* tests. Apelin administration significantly attenuated ALO AIMs scores compared to the control group. Specifically, Mann-Whitney analyses revealed significant reductions in AIMs scores for session 1 (U=31.00, Z= -2.28, p=0.0224), session 3 (U=32.50, Z= -2.20, p=0.0281), and Session 4 (U=14.00, Z= -3.28, p=0.0010). A similar suppressive trend was observed during Session 2 (U=37.00, Z= -1.93, p= 0.05). **F-G.** Graph shows 20-minute time bins for the difference between ALO AIMs of sessions 1 and 4 for control vs apelin groups respectively. **H. I,** and **J** shows the graph for 5-minute bins for days 1, 5 and 9 respectively indicating the progression in the sensitization of the rotational behavior in the control group compared with the apelin treatment groups. The difference was prominent in the later days (day 5, p<0.0001 and day 9, p=0.0019 with time bins x treatment as factors as measured by repeated measure ANOVA). Figure 6B was made using Biorender for publication (License agreement number BI2AGE45BZ).

### Intracellular signaling studies in hemiparkinsonian mice

The molecular mechanisms through which apelin peptide administration attenuates D_1_R hypersensitization and LID were determined by analyzing striatal protein expression and intracellular signaling pathways in hemiparkinsonian mice (Figure 7). Immunoblotting was performed on striatal tissue harvested from intact and lesioned hemispheres following treatment with control liposomes or liposomal apelin13 (Figure 7A). Analysis of desensitization and striatal plasticity markers revealed that apelin13 treatment restored arrestin isoform balance within the lesioned striatum. While absolute levels of arrestin2 and arrestin3 remained relatively comparable across groups (Figure 7B-C), evaluating the percentage increase in the lesioned hemisphere relative to the intact side demonstrated that apelin13 significantly attenuated the excessive accumulation of arrestin3 (p=0.0026) without significantly altering arrestin2 (p=0.312) (Figure 7D). 6-OHDA lesioning and chronic L-DOPA treatment produced marked elevations in phosphorylated ERK (p<0.0001) (Figure 7A and E, G) and phosphorylated p38 (p=0.0019) (Figure 7A and F, G) in the lesioned striatum and intranasal apelin13 had no effect on phosphorylated ERK and p38 compared to control group. Phosphorylated SAPK/JNK isoforms (p46 and p54) showed no significant alterations between control and apelin13 groups (Figure 7A and H, I). Assessment of the Akt mTOR signaling pathway revealed selective modulation of downstream effector targets (Figure 7A, J-L). Phosphorylated Akt at Thr308 (Figure 7J) and Ser473 (Figure 7K), as well as PDK1 phosphorylation (Figure 7M, O), showed no significant differences between control and apelin-13 groups. However, apelin13 treatment significantly reduced hyper activated phospho-mTOR levels in the lesioned striatum (p=0.0446) (Figure 7N, O). The parkin expression remained unaltered (Figure 7A, P), whereas there was reduced α-synuclein levels (p=0.044) (Figure 7A, Q) in the lesioned hemisphere of the apelin peptide treatment group. Reduced striatal ΔFosB levels and attenuated lesion-associated increase in arrestin-3 (p=0.0426) (Figure 7A, R) indicates a targeted dampening of downstream translational control cascades with apelin peptide treatment.

**Figure 7:**
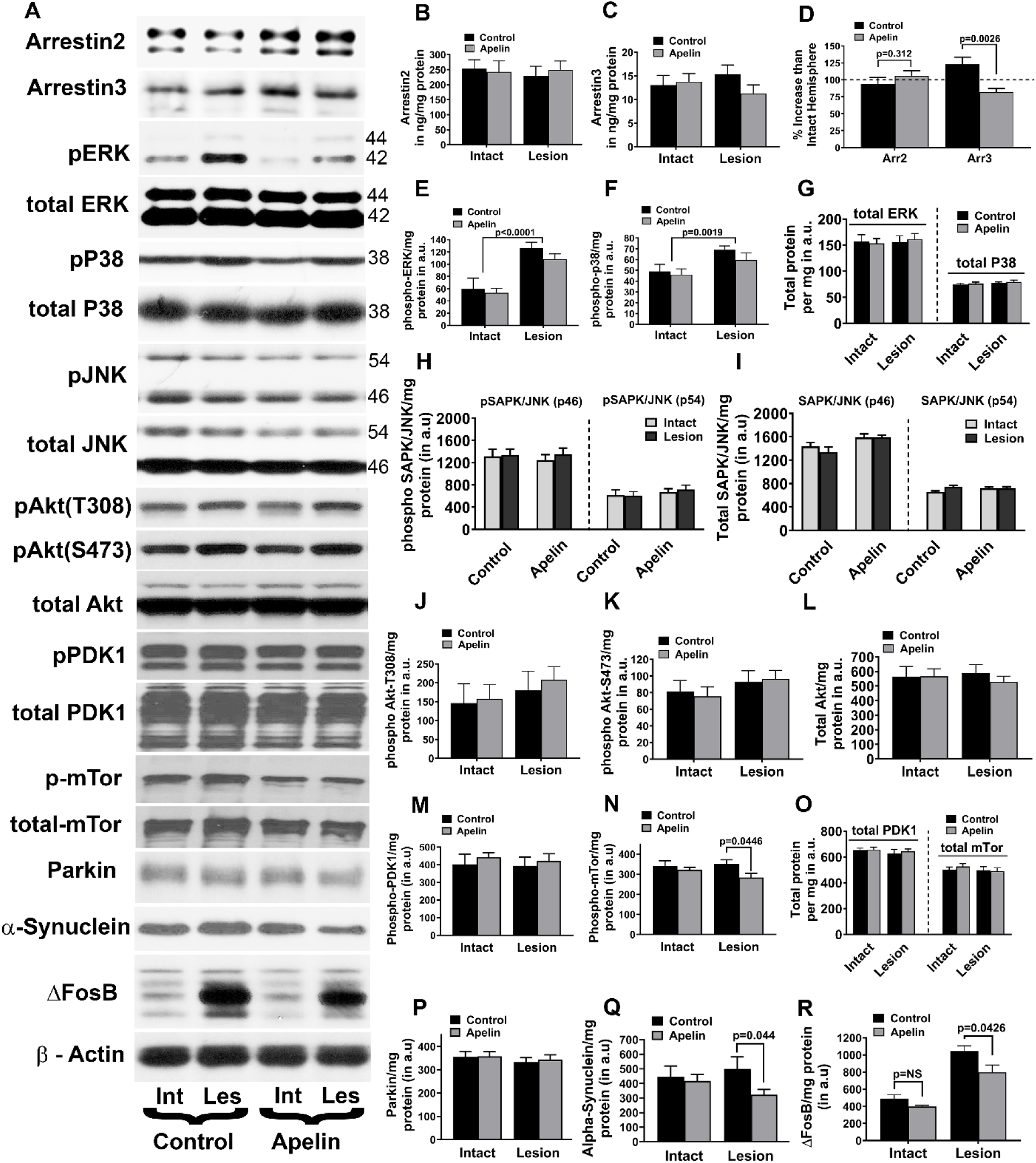
**A.** Western blot pictures showing protein expression. The representative samples for arrestin2, arrestin3, activated (phospho) and total proteins for ERK, p38, SAPK/JNK, Akt (Thr308, Ser473), PDK1, mTor, Parkin, αSynuclein and ΔFosB are shown from the intact and lesioned hemispheres of mice treated with blank control and apelin liposomal formulations in hemiparkinsonian mice model. βactin was used as a loading control for the samples. **B.** Graphical representation of arrestin2 expression, **C.** Graphical representation of arrestin3 expression, **D.** comparison of percentage increase in the expression of arrestin 2 and 3 in the lesioned hemisphere from that of the intact side in control vs apelin peptide treatment groups respectively. **E.** Graphical representation of phospho-ERK expression, **F.** phospho-p38 expression and **G** shows the levels of total ERK and p38 proteins. **H-I.** Graphical representation for the expression of phospho and total SAPK/JNK respectively. **J, K** and **L.** Graphical representation for the expression of phospho Akt (Thr308), (Ser473) and total Akt respectively. **M, N** and **O.** Graphical representation for the expression of phospho PDK1, mTor and their total proteins respectively. **P, Q** and **R** shows the graphical representation of parkin (P), αSynuclein (Q) and ΔFosB (R) respectively.

#### Pathway specific approach using apelin peptide to ameliorate LID

The pathway specific contribution of apelin receptors in regulating LID was determined by evaluating the spatial expression of apelin receptors across striatal cell types and selective cre recombinase driven viral expression in direct or indirect pathway medium spiny neurons (Figure 8). Immunofluorescence staining demonstrated robust expression of APJR in TH positive dopaminergic terminals within the striatum and substantia nigra pars compacta (Figure 8A-F). Co-localization analysis confirmed that apelin receptors are expressed in direct pathway medium spiny neurons labeled in Drd1a-cre mice (Figure 8G) as well as indirect pathway medium spiny neurons labeled in Adora2a-cre mice (Figure 8H). To selectively express apelin36 within specific striatal projection pathways, hemiparkinsonian Drd1a-cre or Adora2a-cre mice were infused with cre dependent double inverted open reading frame adeno-associated virus vectors (AAV-DIO) into the lesioned striatum (Figure 8I-N). Unilaterally lesioned Drd1a-cre mice pretested with daily L-DOPA (25mg/Kg) displayed baseline rotational responses (Figure 8I). Following viral delivery and surgical recovery, Drd1a-cre control mice expressing DIO-GFP or DIO scrambled apelin-36 exhibited prominent motor sensitization with increased contralateral rotations during 10 days of post-AAV L-DOPA testing (p<0.0001) (Figure 8I). Conversely, selective targeted expression of DIO apelin36 within direct pathway medium spiny neurons completely prevented this motor sensitization, maintaining stable low contralateral rotations throughout the testing period (p<0.0001) (Figure 8I). Confocal imaging and immunoblotting verified successful cre recombinase dependent expression of GFP co-cistronic with apelin36 or scrambled apelin36 in cre positive direct pathway neurons (Figure 8J, K). In contrast, targeted viral expression of apelin36 within indirect pathway medium spiny neurons of Adora2a-cre hemiparkinsonian mice failed to modify dyskinetic behavior (Figure 8L). Following 10 days of post-AAV L-DOPA administration, Adora2a-cre mice expressing DIO apelin36 showed escalating contralateral rotations that were statistically indistinguishable from control mice expressing DIO-GFP or DIO scrambled apelin36 (p=NS) (Figure 8L). Confocal microscopy and immunoblotting confirmed equivalent cre co-localization and viral transduction across all Adora2a-cre experimental groups (Figure 8M, N). TH expression in the immunoblots shown in figures 8K and N shows complete loss of dopaminergic projections in the lesioned hemispheres.

**Figure 8:**
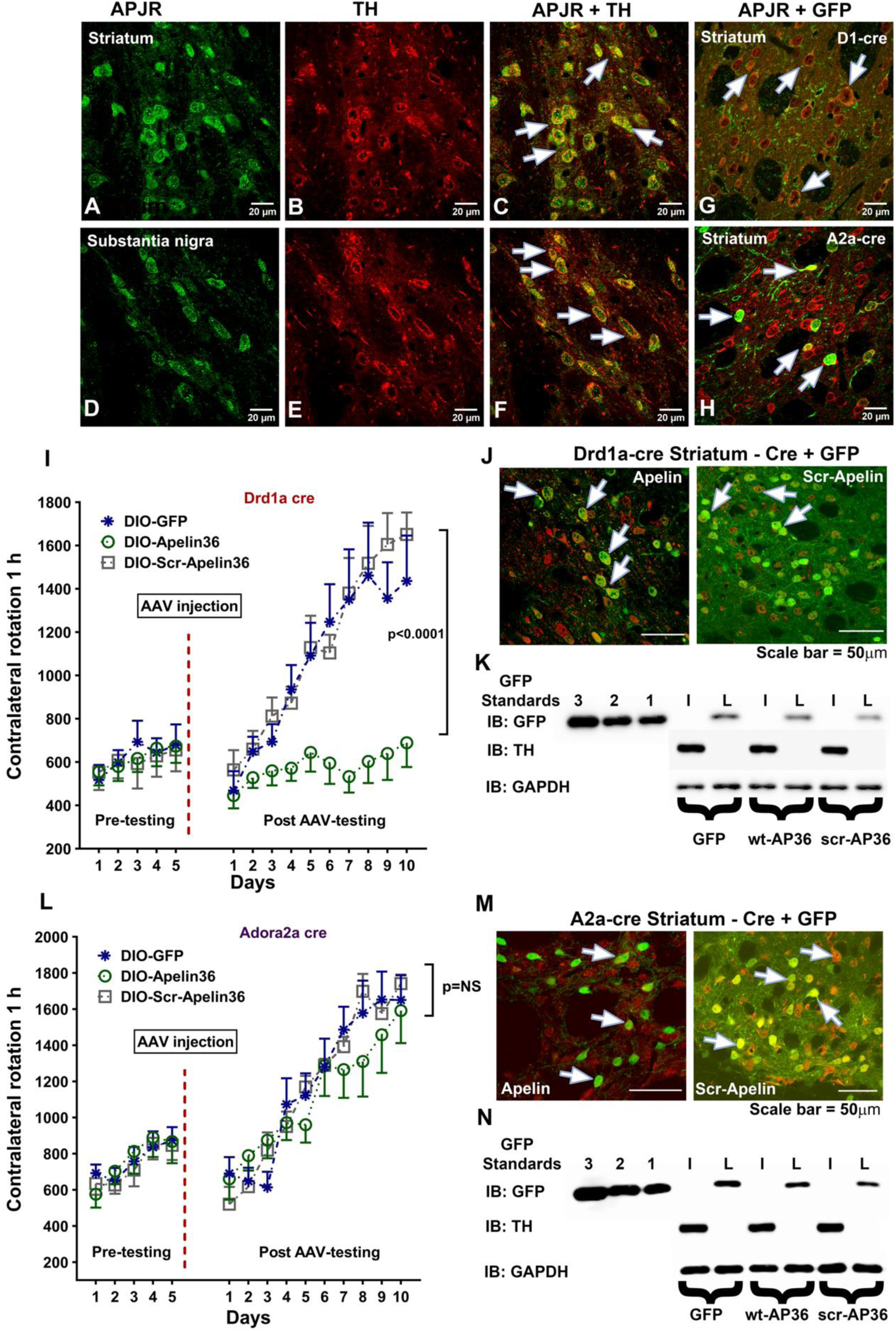
Targeting the dopaminergic direct and indirect pathway neurons to identify the neuronal circuit responsible for the action of APJR in LID. Panels **A-B** show the expression of APJR (green) in TH positive neurons (red) in striatum. Panel **C** shows the merged signal (white arrows) for APJR and TH co-expression. Panels **D-E** show the expression of APJR (green) in TH positive neurons (red) in substantia nigra. Panel **F** shows the merged signal (white arrows) for APJR and TH co-expression. **G-H** panels show the expression of APJR in D1 and A2a-cre positive neurons respectively. The cre positive neurons were labeled with GFP expressed via AAV-DIO-GFP. The expression of APJR was visualized co-expressing in cre positive GFP expressing neuronal medium spiny neurons in the striatum. **I.** Contralateral rotational studies in hemiparkinsonian mouse model with double inverted open reading frame (DIO) AAV mediated expression of apelin36 and scrambled apelin36 peptide with AAV-CAG-DIO-GFP injected mice serving as positive control. The apelin36 constructs had co-cistronic expression of GFP. D1cre mice unilaterally lesioned with 6-OHDA were chronically treated with L-DOPA (25mg/Kg) for 5 days to measure rotational behavior and were grouped for AAV injection into the ipsilateral side of the lesion and post tested after 4 weeks of surgical recovery with L-DOPA for another 10 days. The increase in contralateral rotational behavior for the experimental groups is shown. The cell type-specific therapeutic effects of Apelin overexpression on rotational asymmetry, contralateral rotations (1h) were assessed across pre-testing and 10 post-injection days in D1Cre mice overexpressing DIO-wt-Apelin36 (n=8), DIO-scr-Apelin36 (n=8), or GFP controls (n=7), animals that died during the course of the study were excluded from the statistical analysis. A two-way repeated measures ANOVA revealed a significant main effect of Genotype (F_2,180_ = 11.47, p=0.0005), a significant main effect of Days (p<0.0001), and a significant Genotype × Days interaction (F_18,180_=4.68, p<0.0001). Bonferroni’s post-hoc analyses across days demonstrated that wild-type Apelin36 (wt-Apelin36) overexpression significantly attenuated contralateral rotational behavior compared to both GFP controls (p<0.0001) and the scrambled control group (scrApelin36; p<0.0001). No significant difference in rotational behavior was detected between the GFP and scrAPLN36 groups (p=0.3629). **J.** Immunostaining for GFP and cre in AAV-df-apelin36 and AAV-df-scrambled-apelin36 injected D1cre mice line. The AAV was injected ipsilateral to the side of the lesion. The white arrows show the co-expression of GFP (green) signal in cre (red) positive neurons in D1cre mice. **K.** Western blot pictures showing GFP expression in control GFP and experimental apelin36 and scramble apelin36 groups. **L.** Contralateral rotational studies in hemiparkinsonian mouse model with AAV-DIO mediated expression of apelin36 and scrambled apelin36 peptide with AAV-CAG-DIO-GFP injected mice serving as positive control. The apelin36 constructs had co-cistronic expression of GFP. A2a-cre mice unilaterally lesioned with 6-OHDA were chronically treated with L-DOPA (25mg/Kg) for 5 days to measure rotational behavior and were grouped for AAV injection into the ipsilateral side of the lesion and post tested after 4 weeks of surgical recovery with L-DOPA for another 10 days. There was no difference in the contralateral rotational behavior tested with either apelin36 or scrambled 36 overexpression in A2a (D2R) cre lines compared with the GFP control. **M.** Immunostaining for GFP and cre in AAV-DIO-apelin36 and AAV-DIO-scrambled-apelin36 injected A2a-cre mice line. The AAV was injected ipsilateral to the side of the lesion. The white arrows show the co-expression of GFP (green) signal in cre (red) positive neurons in A2a-cre mice. **N.** Western blot pictures showing GFP expression in control GFP and experimental apelin36 and scramble apelin36 groups.

## Discussion

Loss of nigrostriatal dopamine transporters combined with non-physiological dopamine synthesis induces persistent hyperactivation of direct pathway MSNs leading to aberrant striatal plasticity and over inhibition of basal ganglia output structures, manifesting clinically as AIMs^6,16,48–50^. Targeting GPCR signaling networks within direct pathway MSNs represents an attractive approach to reestablish physiological signaling without compromising the antiparkinsonian benefits of L-DOPA. APJR are co-expressed with D_1_ and D_2_R in MSNs and targeting the apelin/APJR axis offers a specific avenue to modulate direct pathway output. The APJ receptor, a class A GPCR, shares significant sequence homology with the angiotensin II receptor type 1 (AT_1_R) and is coupled to multiple G proteins, including Gα_i_ and Gα_q32_. Apelin, the endogenous ligand for the APJR, is produced as a 77-amino acid pre-proapelin and processed into active fragments, including apelin-13, apelin-17, and apelin-36^51–53^. These peptides have been implicated in various physiological processes, including cardiovascular regulation, fluid homeostasis, and energy metabolism^30,54–58^. In the CNS, the APJR and apelin are expressed in regions such as the basal ganglia, which are critical for motor control^55,56^. Emerging evidence suggests that the APJR-apelin system may play a role in modulating neuronal excitability and provide neuroprotection ^28,59,60^. Studies have demonstrated that apelin exerts neuroprotective effects on dopaminergic neurons in the substantia nigra pars compacta, the primary site of neurodegeneration in PD^61^. Despite these promising findings, the role of the APJ receptor and apelin in the context of LID has not been explored. Activation of APJR by apelin peptide treatment engages alternative intracellular signaling pathways, such as recruitment of G_αi_ or arrestin, which can cross regulate hyperactive D_1_R signaling cascades.

The cAMP assays demonstrate the delicate receptor cross talk and signaling balance that exists between G_αs_ coupled and G_αi/o_ coupled GPCRs. Under baseline conditions, DA-D_2_R activation acts as an intracellular brake on DA-D_1_R mediated AC activation ^62–64^, dampening cAMP synthesis and downstream PKA cascades. When D_2_R activity is compromised, blocked by antagonists, or functionally uncoupled, this inhibitory brake is removed. The resulting uninhibited G_αs_ signaling leads to hyperactivation of cAMP production and excessive stimulation of downstream effectors such as PKA/DARPP-32/MAPK pathway proteins in striatal direct pathway MSNs^65–68^. This mechanism provides direct insight into the intracellular pathology of LID. In Parkinsonian states, denervation and chronic non-physiological L-DOPA administration sensitize D_1_R on direct pathway neurons^69–71^. Unchecked D_1_R signaling drives pathological cAMP accumulation and abnormal transcriptional plasticity^72–74^, resulting in involuntary dyskinetic movements^75,76^. Restoring balance to this hyper-active cAMP signaling cascade without directly blocking D1R is critical for therapeutic interventions. Targeting the APJR, a G_αi/o_ coupled receptor expressed alongside DA-D_1_R in direct pathway medium spiny neurons, offers a compelling strategy to restore signaling homeostasis. Because APJR activation engages G_αi/o_ pathways and internalizes via arrestin recruitment, apelin peptide treatment can directly counterbalance DA-D1R hyper-sensitization. By introducing a targeted G_αi/o_ inhibitory signal specifically within direct pathway neurons, apelin treatment can suppress aberrant AC activity and reduce excessive cAMP production.

The quantitative kinetics and cellular imaging data demonstrate that DA-D_1_R undergo classical homologous desensitization via rapid arrestin2 and arrestin3 recruitment following agonist exposure. Under physiological conditions, this arrestin mediated recruitment serves as an essential negative feedback loop^77–84^, uncoupling the receptor from G_αs_ proteins, internalizing the receptor complex, and preventing prolonged intracellular cyclic AMP pathway overactivation^81,82^. Impairments or saturation of this homologous desensitization machinery contribute significantly to the pathophysiology of LID^85^. In Parkinsonian models, chronic pulsatile L-DOPA treatment leads to aberrant DA-D_1_R hypersensitization and also drives the expression of arrestin3^86,87^. This maladaptation leaves D_1_R in a persistently active state, with cAMP accumulation and arrestin3 mediated signaling to drive severe dyskinetic motor behaviors^86^. Therapeutic targeting of the APJR provides a mechanism to restore proper receptor desensitization dynamics within striatal direct pathway neurons. Because APJR are co-expressed with D_1_R on direct pathway MSNs, apelin peptide treatment can engage robust G_αi/o_ signaling and arrestin recruitment cascades. APJR activation promotes receptor co-internalization or heterologous desensitization of sensitized D1R complexes^88–92^, efficiently dampening hyperactive cAMP cascades and normalizing MSN excitability to ameliorate LID.

The biochemical and behavioral data establishes that APJR activation by apelin13 or apelin36 exerts strong competitive inhibition over D_1_R mediated signaling pathways. The maladaptive biochemical cascades driven by DA depletion combined with chronic L-DOPA administration leads to persistent D_1_R signaling pathways on striatal direct MSNs. This unbuffered signaling drives abnormal overactivation of the MAPK pathways, leading to nuclear translocation and long term accumulation of the stable transcription factor ΔFosB. Accumulated ΔFosB mediates aberrant gene expression and transcriptional plasticity that perpetuate dyskinetic motor behaviors ^47,93–95^. By engaging G_αi/o_ signaling cascades, APJR activation directly antagonizes AC activity, blunting D_1_R driven cAMP synthesis and downstream PKA stimulation. Furthermore, apelin treatment sequesters intracellular arrestin pools away from D_1_R. This competitive engagement prevents aberrant D_1_R signaling cascades while driving APJR endocytosis. These findings hold direct relevance for modulating direct pathway striatal MSNs in LID. By dampening excessive cAMP production and restoring intracellular signaling homeostasis without directly blocking D_1_R, APJR targeting represents a novel therapeutic strategy to ameliorate LID while preserving motor function. The viral vector mediated expression of apelin36 in the striatum effectively prevents the behavioral sensitization characteristic of LID. Striatal transduction of apelin36 directly into FoxP positive medium spiny neurons halts this pathological motor sensitization. The endogenous production of apelin36 provides continuous G_αi/o_ signaling tone and facilitates receptor desensitization pathways. This intracellular modulation to attenuate hyperactive D1R responses indirectly without compromising the basic prokinetic drive required to alleviate Parkinsonian akinesia confirm that targeting the striatal apelin/APJ receptor axis in vivo provides a potent non-dopaminergic therapeutic strategy to ameliorate LID in PD.

Furthermore, the selective downregulation of phospho-mTOR by apelin highlights its role in normalizing local translational control associated with dyskinetic synaptic plasticity^96,97^. Apelin-13 reduced mTOR phosphorylation in the lesioned striatum, whereas ERK and p38 phosphorylation were not significantly affected. These biochemical findings demonstrate that targeting the striatal APJR provides a robust molecular mechanism to counteract D_1_R hypersensitization and alleviate LID in PD. Targeting the striatal APJR effectively intercepts this pathological cascade at multiple nodes.

The pathway specific findings identify direct pathway medium spiny neurons as the essential anatomical and functional locus where apelin receptor activation acts to ameliorate LID. Although apelin receptors are expressed across both direct and indirect striatal projection neurons, as well as presynaptic dopaminergic terminals, targeted manipulation reveals a selective vulnerability and therapeutic window exclusive to the direct pathway. Cre dependent expression of apelin36 specifically in Drd1a-cre neurons restores intracellular signaling homeostasis within these direct pathway MSNs. By engaging G_αi/o_ signaling and arrestin mediated desensitization cascades directly in Drd1a expressing cells, apelin36 suppresses hyperactive AC activity and prevents behavioral sensitization. Conversely, apelin36 expression within indirect pathway medium spiny neurons in Adora2a-cre mice provided no protection against LID. Because dyskinetic sensitization is primarily driven by D_1_R hyper-activation in direct pathway projection neurons, modulating G_α/i_ tone in indirect pathway neurons is insufficient to counteract the direct pathway output imbalance. These results prove that the therapeutic anti-dyskinetic actions of apelin peptide treatment depend specifically on targeting the APJR axis in direct pathway striatal MSNs. Apelin peptide treatment targeting striatal APJR attenuates aberrant cAMP production and downstream signaling and pharmacological targeting of the APJR provides a potential strategy to normalize direct pathway output and alleviate LID in PD.

## Author contributions

MRA and JR conceived the study plan. MRA designed, executed, and wrote the manuscript with input from co-authors. CZ and MRA designed the cAMP and nanoBiT assays with receptor signaling in cell lines. MI and MRA did the HPLC purification of the apelin13 peptide. Animal surgeries, behavioral and biochemical studies were performed by MRA. CZ, MSA, MM, MMB and MI contributed to data curation, tissue collection, and western blot analysis. JR supervised the overall study. The authors declare no conflict of interest.

## Acknowledgements

The authors thank Drs. Eugenia Gurevich and Vsevolod Gurevich of Department of Pharmacology, Vanderbilt University for their help with the resources. The authors thank Dr. Tina Iverson of Department of Pharmacology for instrumental resources related to cAMP and nanoBiT assays. The confocal microscopic imaging was performed in part through the use of the Vanderbilt Cell Imaging Shared Resource (supported by NIH grants CA68485, DK20593, DK58404, DK59637 and EY08126). The authors thank Dr. Mehrdad Shamloo of Stanford University SBFNL for the shared resources for animal surgeries and behavioral studies. The authors also thank Dr. Mark Schnitzer Lab, Stanford University for AAV-CAG-DIO-Ax vector for cloning and mouse D1 and A2a cre lines for behavioral experiments.

## Data Availability

The raw/processed data required to reproduce these findings cannot be shared at this time due to technical or time limitations. The authors would share the appropriate raw data/information upon reasonable request.

## References

1. Kraus, P.H., Klotz, P., Fischer, A., and Przuntek, H. (1987). Assessment of symptoms of Parkinson’s disease by apparative methods. J Neural Transm Suppl 25, 89–96.

2. Lee, R.G. (1987). Physiology of the basal ganglia and pathophysiology of Parkinson’s disease. Can J Neurol Sci 14, 373–380. 10.1017/s0317167100037768.

3. Cools, A.R. (1984). Basal ganglia and Parkinson’s disease: neurobiological and pharmacological aspects in animals and man. Clin Neurol Neurosurg 86, 178–195. 10.1016/0303-8467(84)90195-1.

4. Marsden, C.D. (1984). Function of the basal ganglia as revealed by cognitive and motor disorders in Parkinson’s disease. Can J Neurol Sci 11, 129–135. 10.1017/s031716710004628x.

5. Zamir, N., Skofitsch, G., Bannon, M.J., Helke, C.J., Kopin, I.J., and Jacobowitz, D.M. (1984). Primate model of Parkinson’s disease: alterations in multiple opioid systems in the basal ganglia. Brain Res 322, 356–360. 10.1016/0006-8993(84)90133-1.

6. Cenci, M.A., and Lindgren, H.S. (2007). Advances in understanding L-DOPA-induced dyskinesia. Curr Opin Neurobiol 17, 665–671. 10.1016/j.conb.2008.01.004.

7. Brotchie, J.M. (1999). Advances in understanding the neural mechanisms underlying L-DOPA-induced dyskinesia. Adv Neurol 80, 71–85.

8. Winkler, C., Kirik, D., Bjorklund, A., and Cenci, M.A. (2002). L-DOPA-induced dyskinesia in the intrastriatal 6-hydroxydopamine model of parkinson’s disease: relation to motor and cellular parameters of nigrostriatal function. Neurobiol Dis 10, 165–186. 10.1006/nbdi.2002.0499.

9. Garcia-Escrig, M., and Bermejo-Pareja, F. (1999). [Motor complications of treatment with levodopa in Parkinson’s disease]. Rev Neurol 28, 799–809.

10. Cenci, M.A., Lee, C.S., and Bjorklund, A. (1998). L-DOPA-induced dyskinesia in the rat is associated with striatal overexpression of prodynorphin- and glutamic acid decarboxylase mRNA. Eur J Neurosci 10, 2694–2706.

11. Henry, B., Crossman, A.R., and Brotchie, J.M. (1998). Characterization of a rodent model in which to investigate the molecular and cellular mechanisms underlying the pathophysiology of L-dopa-induced dyskinesia. Adv Neurol 78, 53–61.

12. Bagetta, V., Ghiglieri, V., Sgobio, C., Calabresi, P., and Picconi, B. (2010). Synaptic dysfunction in Parkinson’s disease. Biochem Soc Trans 38, 493–497. 10.1042/BST0380493.

13. Cenci, M.A., and Konradi, C. (2010). Maladaptive striatal plasticity in L-DOPA-induced dyskinesia. Prog Brain Res 183, 209–233. 10.1016/S0079-6123(10)83011-0.

14. Heiman, M., Heilbut, A., Francardo, V., Kulicke, R., Fenster, R.J., Kolaczyk, E.D., Mesirov, J.P., Surmeier, D.J., Cenci, M.A., and Greengard, P. (2014). Molecular adaptations of striatal spiny projection neurons during levodopa-induced dyskinesia. Proc Natl Acad Sci U S A 111, 4578–4583. 10.1073/pnas.1401819111.

15. Kuter, K.Z., Cenci, M.A., and Carta, A.R. (2020). The role of glia in Parkinson’s disease: Emerging concepts and therapeutic applications. Prog Brain Res 252, 131–168. 10.1016/bs.pbr.2020.02.004.

16. Baas, H. (2000). Dyskinesia in Parkinson’s disease. Pathophysiology and clinical risk factors. J Neurol 247 Suppl 4, IV/12-16. 10.1007/pl00007767.

17. Lindgren, H.S., Ohlin, K.E., and Cenci, M.A. (2009). Differential involvement of D1 and D2 dopamine receptors in L-DOPA-induced angiogenic activity in a rat model of Parkinson’s disease. Neuropsychopharmacology 34, 2477–2488. 10.1038/npp.2009.74.

18. Guigoni, C., Aubert, I., Li, Q., Gurevich, V.V., Benovic, J.L., Ferry, S., Mach, U., Stark, H., Leriche, L., Hakansson, K., et al. (2005). Pathogenesis of levodopa-induced dyskinesia: focus on D1 and D3 dopamine receptors. Parkinsonism Relat Disord 11 Suppl 1, S25–29. 10.1016/j.parkreldis.2004.11.005.

19. Bezard, E., Gross, C.E., Qin, L., Gurevich, V.V., Benovic, J.L., and Gurevich, E.V. (2005). L-DOPA reverses the MPTP-induced elevation of the arrestin2 and GRK6 expression and enhanced ERK activation in monkey brain. Neurobiol Dis 18, 323–335. 10.1016/j.nbd.2004.10.005.

20. Bido, S., Solari, N., Indrigo, M., D’Antoni, A., Brambilla, R., Morari, M., and Fasano, S. (2015). Differential involvement of Ras-GRF1 and Ras-GRF2 in L-DOPA-induced dyskinesia. Ann Clin Transl Neurol 2, 662–678. 10.1002/acn3.202.

21. Bonito-Oliva, A., Feyder, M., and Fisone, G. (2011). Deciphering the Actions of Antiparkinsonian and Antipsychotic Drugs on cAMP/DARPP-32 Signaling. Front Neuroanat 5, 38. 10.3389/fnana.2011.00038.

22. Cerovic, M., d’Isa, R., Tonini, R., and Brambilla, R. (2013). Molecular and cellular mechanisms of dopamine-mediated behavioral plasticity in the striatum. Neurobiol Learn Mem 105, 63–80. 10.1016/j.nlm.2013.06.013.

23. Feyder, M., Bonito-Oliva, A., and Fisone, G. (2011). L-DOPA-Induced Dyskinesia and Abnormal Signaling in Striatal Medium Spiny Neurons: Focus on Dopamine D1 Receptor-Mediated Transmission. Front Behav Neurosci 5, 71. 10.3389/fnbeh.2011.00071.

24. Carriba, P., Ortiz, O., Patkar, K., Justinova, Z., Stroik, J., Themann, A., Muller, C., Woods, A.S., Hope, B.T., Ciruela, F., et al. (2007). Striatal adenosine A2A and cannabinoid CB1 receptors form functional heteromeric complexes that mediate the motor effects of cannabinoids. Neuropsychopharmacology 32, 2249–2259. 10.1038/sj.npp.1301375.

25. Ferre, S., Lluis, C., Justinova, Z., Quiroz, C., Orru, M., Navarro, G., Canela, E.I., Franco, R., and Goldberg, S.R. (2010). Adenosine-cannabinoid receptor interactions. Implications for striatal function. Br J Pharmacol 160, 443–453. 10.1111/j.1476-5381.2010.00723.x.

26. Stockwell, J., Jakova, E., and Cayabyab, F.S. (2017). Adenosine A1 and A2A Receptors in the Brain: Current Research and Their Role in Neurodegeneration. Molecules 22. 10.3390/molecules22040676.

27. Ivanov, M.N., Stoyanov, D.S., Pavlov, S.P., and Tonchev, A.B. (2022). Distribution, Function, and Expression of the Apelinergic System in the Healthy and Diseased Mammalian Brain. Genes (Basel) 13. 10.3390/genes13112172.

28. Luo, H., Han, L., and Xu, J. (2020). Apelin/APJ system: A novel promising target for neurodegenerative diseases. J Cell Physiol 235, 638–657. 10.1002/jcp.29001.

29. O’Donnell, L.A., Agrawal, A., Sabnekar, P., Dichter, M.A., Lynch, D.R., and Kolson, D.L. (2007). Apelin, an endogenous neuronal peptide, protects hippocampal neurons against excitotoxic injury. J Neurochem 102, 1905–1917. 10.1111/j.1471-4159.2007.04645.x.

30. Reaux, A., Gallatz, K., Palkovits, M., and Llorens-Cortes, C. (2002). Distribution of apelin-synthesizing neurons in the adult rat brain. Neuroscience 113, 653–662. 10.1016/s0306-4522(02)00192-6.

31. O’Dowd, B.F., Heiber, M., Chan, A., Heng, H.H., Tsui, L.C., Kennedy, J.L., Shi, X., Petronis, A., George, S.R., and Nguyen, T. (1993). A human gene that shows identity with the gene encoding the angiotensin receptor is located on chromosome 11. Gene 136, 355–360. 10.1016/0378-1119(93)90495-o.

32. Pitkin, S.L., Maguire, J.J., Bonner, T.I., and Davenport, A.P. (2010). International Union of Basic and Clinical Pharmacology. LXXIV. Apelin receptor nomenclature, distribution, pharmacology, and function. Pharmacol Rev 62, 331–342. 10.1124/pr.110.002949.

33. Hu, S., Wang, D., Liu, W., Wang, Y., Chen, J., and Cai, X. (2024). Apelin receptor dimer: Classification, future prospects, and pathophysiological perspectives. Biochim Biophys Acta Mol Basis Dis 1870, 167257. 10.1016/j.bbadis.2024.167257.

34. Goupil, E., Laporte, S.A., and Hebert, T.E. (2013). GPCR heterodimers: asymmetries in ligand binding and signalling output offer new targets for drug discovery. Br J Pharmacol 168, 1101–1103. 10.1111/bph.12040.

35. Bai, B., Cai, X., Jiang, Y., Karteris, E., and Chen, J. (2014). Heterodimerization of apelin receptor and neurotensin receptor 1 induces phosphorylation of ERK(1/2) and cell proliferation via Galphaq-mediated mechanism. J Cell Mol Med 18, 2071–2081. 10.1111/jcmm.12404.

36. Bai, B., Liu, L., Zhang, N., Wang, C., Jiang, Y., and Chen, J. (2014). Heterodimerization of human apelin and bradykinin 1 receptors: novel signal transduction characteristics. Cell Signal 26, 1549–1559. 10.1016/j.cellsig.2014.03.022.

37. Li, Y., Chen, J., Bai, B., Du, H., Liu, Y., and Liu, H. (2012). Heterodimerization of human apelin and kappa opioid receptors: roles in signal transduction. Cell Signal 24, 991–1001. 10.1016/j.cellsig.2011.12.012.

38. Chen, J., Wang, Z., Zhang, R., Yin, H., Wang, P., Wang, C., and Jiang, Y. (2023). Heterodimerization of apelin and opioid receptor-like 1 receptors mediates apelin-13-induced G protein biased signaling. Life Sci 328, 121892. 10.1016/j.lfs.2023.121892.

39. Wan, L., Xu, F., Liu, C., Ji, B., Zhang, R., Wang, P., Wu, F., Pan, Y., Yang, C., Wang, C., and Chen, J. (2020). Transmembrane peptide 4 and 5 of APJ are essential for its heterodimerization with OX1R. Biochem Biophys Res Commun 521, 408–413. 10.1016/j.bbrc.2019.10.146.

40. Siddiquee, K., Hampton, J., McAnally, D., May, L., and Smith, L. (2013). The apelin receptor inhibits the angiotensin II type 1 receptor via allosteric trans-inhibition. Br J Pharmacol 168, 1104–1117. 10.1111/j.1476-5381.2012.02192.x.

41. Siddiquee, K., Hampton, J., Khan, S., Zadory, D., Gleaves, L., Vaughan, D.E., and Smith, L.H. (2011). Apelin protects against angiotensin II-induced cardiovascular fibrosis and decreases plasminogen activator inhibitor type-1 production. J Hypertens 29, 724–731. 10.1097/HJH.0b013e32834347de.

42. Dahl, L., Kotliar, I.B., Bendes, A., Dodig-Crnkovic, T., Fromm, S., Elofsson, A., Uhlen, M., Sakmar, T.P., and Schwenk, J.M. (2023). Multiplexed selectivity screening of anti-GPCR antibodies. Sci Adv 9, eadf9297. 10.1126/sciadv.adf9297.

43. Zheng, C., Javitch, J.A., Lambert, N.A., Donthamsetti, P., and Gurevich, V.V. (2023). In-Cell Arrestin-Receptor Interaction Assays. Curr Protoc 3, e890. 10.1002/cpz1.890.

44. Ahmed, M.R., Inayathullah, M., Morton, M., Pothineni, V.R., Kim, K., Ahmed, M.S., Babar, M.M., and Rajadas, J. (2024). Intranasal delivery of liposome encapsulated flavonoids ameliorates l-DOPA induced dyskinesia in hemiparkinsonian mice. Biomaterials 311, 122680. 10.1016/j.biomaterials.2024.122680.

45. Bochkov, Y.A., and Palmenberg, A.C. (2006). Translational efficiency of EMCV IRES in bicistronic vectors is dependent upon IRES sequence and gene location. Biotechniques 41, 283–284, 286, 288 passim. 10.2144/000112243.

46. Ahmed, M.R., Zheng, C., Dunning, J.L., Ahmed, M.S., Ge, C., Pair, F.S., Gurevich, V.V., and Gurevich, E.V. (2024). Arrestin-3-assisted activation of JNK3 mediates dopaminergic behavioral sensitization. Cell Rep Med 5, 101623. 10.1016/j.xcrm.2024.101623.

47. Ahmed, M.R., Jayakumar, M., Ahmed, M.S., Zamaleeva, A.I., Tao, J., Li, E.H., Job, J.K., Pittenger, C., Ohtsu, H., and Rajadas, J. (2019). Pharmacological antagonism of histamine H2R ameliorated L-DOPA-induced dyskinesia via normalization of GRK3 and by suppressing FosB and ERK in PD. Neurobiol Aging 81, 177–189. 10.1016/j.neurobiolaging.2019.06.004.

48. Carta, M., Lindgren, H.S., Lundblad, M., Stancampiano, R., Fadda, F., and Cenci, M.A. (2006). Role of striatal L-DOPA in the production of dyskinesia in 6-hydroxydopamine lesioned rats. J Neurochem 96, 1718–1727. 10.1111/j.1471-4159.2006.03696.x.

49. Sugiyama, K., Kuroiwa, M., Shuto, T., Hwang, S., Oh, Y.S., and Nishi, A. (2025). Subregion-specific suppression of dopamine D1 receptor expression prevents L-DOPA-induced dyskinesia in a mouse model of Parkinson’s disease. Neurosci Lett 859-861, 138273. 10.1016/j.neulet.2025.138273.

50. Sugiyama, K., Kuroiwa, M., Shuto, T., Ohnishi, Y.N., Kawahara, Y., Miyamoto, Y., Fukuda, T., and Nishi, A. (2021). Subregion-Specific Regulation of Dopamine D1 Receptor Signaling in the Striatum: Implication for L-DOPA-Induced Dyskinesia. J Neurosci 41, 6388–6414. 10.1523/JNEUROSCI.0373-21.2021.

51. Tatemoto, K., Hosoya, M., Habata, Y., Fujii, R., Kakegawa, T., Zou, M.X., Kawamata, Y., Fukusumi, S., Hinuma, S., Kitada, C., et al. (1998). Isolation and characterization of a novel endogenous peptide ligand for the human APJ receptor. Biochem Biophys Res Commun 251, 471–476. 10.1006/bbrc.1998.9489.

52. Masri, B., Knibiehler, B., and Audigier, Y. (2005). Apelin signalling: a promising pathway from cloning to pharmacology. Cell Signal 17, 415–426. 10.1016/j.cellsig.2004.09.018.

53. De Mota, N., Reaux-Le Goazigo, A., El Messari, S., Chartrel, N., Roesch, D., Dujardin, C., Kordon, C., Vaudry, H., Moos, F., and Llorens-Cortes, C. (2004). Apelin, a potent diuretic neuropeptide counteracting vasopressin actions through inhibition of vasopressin neuron activity and vasopressin release. Proc Natl Acad Sci U S A 101, 10464–10469. 10.1073/pnas.0403518101.

54. Han, R.W., Xu, H.J., Zhang, R.S., and Wang, R. (2014). The role of apelin-13 in novel object recognition memory. Peptides 62, 155–158. 10.1016/j.peptides.2014.10.003.

55. Matsumoto, M., Hidaka, K., Akiho, H., Tada, S., Okada, M., and Yamaguchi, T. (1996). Low stringency hybridization study of the dopamine D4 receptor revealed D4-like mRNA distribution of the orphan seven-transmembrane receptor, APJ, in human brain. Neurosci Lett 219, 119–122. 10.1016/s0304-3940(96)13198-0.

56. Matsumoto, M., Hidaka, K., Tada, S., Tasaki, Y., and Yamaguchi, T. (1996). Low levels of mRNA for dopamine D4 receptor in human cerebral cortex and striatum. J Neurochem 66, 915–919. 10.1046/j.1471-4159.1996.66030915.x.

57. Taheri, S., Murphy, K., Cohen, M., Sujkovic, E., Kennedy, A., Dhillo, W., Dakin, C., Sajedi, A., Ghatei, M., and Bloom, S. (2002). The effects of centrally administered apelin-13 on food intake, water intake and pituitary hormone release in rats. Biochem Biophys Res Commun 291, 1208–1212. 10.1006/bbrc.2002.6575.

58. Zhou, J.X., Shuai, N.N., Wang, B., Jin, X., Kuang, X., and Tian, S.W. (2021). Neuroprotective gain of Apelin/APJ system. Neuropeptides 87, 102131. 10.1016/j.npep.2021.102131.

59. Cheng, B., Chen, J., Bai, B., and Xin, Q. (2012). Neuroprotection of apelin and its signaling pathway. Peptides 37, 171–173. 10.1016/j.peptides.2012.07.012.

60. Esmaeili-Mahani, S., Haghparast, E., Nezhadi, A., Abbasnejad, M., and Sheibani, V. (2021). Apelin-13 prevents hippocampal synaptic plasticity impairment in Parkinsonism rats. J Chem Neuroanat 111, 101884. 10.1016/j.jchemneu.2020.101884.

61. Liu, C., Kang, N.W., Wang, Y., Xue, Y., Chen, X.Y., and Chen, L. (2022). Neuropeptide apelin presented in the dopaminergic neurons modulates the neuronal excitability in the substantia nigra pars compacta. Neuropharmacology 219, 109235. 10.1016/j.neuropharm.2022.109235.

62. Taylor, J.L., Bishop, C., and Walker, P.D. (2005). Dopamine D1 and D2 receptor contributions to L-DOPA-induced dyskinesia in the dopamine-depleted rat. Pharmacol Biochem Behav 81, 887–893. 10.1016/j.pbb.2005.06.013.

63. Farre, D., Munoz, A., Moreno, E., Reyes-Resina, I., Canet-Pons, J., Dopeso-Reyes, I.G., Rico, A.J., Lluis, C., Mallol, J., Navarro, G., et al. (2015). Stronger Dopamine D1 Receptor-Mediated Neurotransmission in Dyskinesia. Mol Neurobiol 52, 1408–1420. 10.1007/s12035-014-8936-x.

64. Rosengarten, H., Schweitzer, J.W., and Friedhoff, A.J. (1986). Selective dopamine D2 receptor reduction enhances a D1-mediated oral dyskinesia in rats. Life Sci 39, 29–35. 10.1016/0024-3205(86)90434-0.

65. Goto, S. (2017). Striatal Galpha(olf)/cAMP Signal-Dependent Mechanism to Generate Levodopa-Induced Dyskinesia in Parkinson’s Disease. Front Cell Neurosci 11, 364. 10.3389/fncel.2017.00364.

66. Santini, E., Sgambato-Faure, V., Li, Q., Savasta, M., Dovero, S., Fisone, G., and Bezard, E. (2010). Distinct changes in cAMP and extracellular signal-regulated protein kinase signalling in L-DOPA-induced dyskinesia. PLoS One 5, e12322. 10.1371/journal.pone.0012322.

67. Santini, E., Valjent, E., Usiello, A., Carta, M., Borgkvist, A., Girault, J.A., Herve, D., Greengard, P., and Fisone, G. (2007). Critical involvement of cAMP/DARPP-32 and extracellular signal-regulated protein kinase signaling in L-DOPA-induced dyskinesia. J Neurosci 27, 6995–7005. 10.1523/JNEUROSCI.0852-07.2007.

68. Hakansson, K., Lindskog, M., Pozzi, L., Usiello, A., and Fisone, G. (2004). DARPP-32 and modulation of cAMP signaling: involvement in motor control and levodopa-induced dyskinesia. Parkinsonism Relat Disord 10, 281–286. 10.1016/j.parkreldis.2004.02.010.

69. Mulas, G., Espa, E., Fenu, S., Spiga, S., Cossu, G., Pillai, E., Carboni, E., Simbula, G., Jadzic, D., Angius, F., et al. (2016). Differential induction of dyskinesia and neuroinflammation by pulsatile versus continuous l-DOPA delivery in the 6-OHDA model of Parkinson’s disease. Exp Neurol 286, 83–92. 10.1016/j.expneurol.2016.09.013.

70. Spigolon, G., and Fisone, G. (2018). Signal transduction in L-DOPA-induced dyskinesia: from receptor sensitization to abnormal gene expression. J Neural Transm (Vienna) 125, 1171–1186. 10.1007/s00702-018-1847-7.

71. Dunnett, S. (2003). L-DOPA, dyskinesia and striatal plasticity. Nat Neurosci 6, 437–438. 10.1038/nn0503-437.

72. Guerra, A., D’Onofrio, V., Asci, F., Ferreri, F., Fabbrini, G., Berardelli, A., and Bologna, M. (2023). Assessing the interaction between L-dopa and gamma-transcranial alternating current stimulation effects on primary motor cortex plasticity in Parkinson’s disease. Eur J Neurosci 57, 201–212. 10.1111/ejn.15867.

73. Bove, F., and Calabresi, P. (2022). Plasticity, genetics, and epigenetics in l-dopa-induced dyskinesias. Handb Clin Neurol 184, 167–184. 10.1016/B978-0-12-819410-2.00009-6.

74. Borgkvist, A., Lieberman, O.J., and Sulzer, D. (2018). Synaptic plasticity may underlie l-DOPA induced dyskinesia. Curr Opin Neurobiol 48, 71–78. 10.1016/j.conb.2017.10.021.

75. Wang, Q., and Zhang, W. (2016). Maladaptive Synaptic Plasticity in L-DOPA-Induced Dyskinesia. Front Neural Circuits 10, 105. 10.3389/fncir.2016.00105.

76. Fieblinger, T., Graves, S.M., Sebel, L.E., Alcacer, C., Plotkin, J.L., Gertler, T.S., Chan, C.S., Heiman, M., Greengard, P., Cenci, M.A., and Surmeier, D.J. (2014). Cell type-specific plasticity of striatal projection neurons in parkinsonism and L-DOPA-induced dyskinesia. Nat Commun 5, 5316. 10.1038/ncomms6316.

77. Skinbjerg, M., Ariano, M.A., Thorsell, A., Heilig, M., Halldin, C., Innis, R.B., and Sibley, D.R. (2009). Arrestin3 mediates D(2) dopamine receptor internalization. Synapse 63, 621–624. 10.1002/syn.20636.

78. Macey, T.A., Liu, Y., Gurevich, V.V., and Neve, K.A. (2005). Dopamine D1 receptor interaction with arrestin3 in neostriatal neurons. J Neurochem 93, 128–134. 10.1111/j.1471-4159.2004.02998.x.

79. Gray, D.L., Allen, J.A., Mente, S., O’Connor, R.E., DeMarco, G.J., Efremov, I., Tierney, P., Volfson, D., Davoren, J., Guilmette, E., et al. (2018). Impaired beta-arrestin recruitment and reduced desensitization by non-catechol agonists of the D1 dopamine receptor. Nat Commun 9, 674. 10.1038/s41467-017-02776-7.

80. Gurevich, V.V., Chen, Q., and Gurevich, E.V. (2018). Arrestins: Introducing Signaling Bias Into Multifunctional Proteins. Prog Mol Biol Transl Sci 160, 47–61. 10.1016/bs.pmbts.2018.07.007.

81. Gurevich, V.V., and Gurevich, E.V. (2015). Arrestins: Critical Players in Trafficking of Many GPCRs. Prog Mol Biol Transl Sci 132, 1–14. 10.1016/bs.pmbts.2015.02.010.

82. Gurevich, V.V., and Gurevich, E.V. (2015). Analyzing the roles of multi-functional proteins in cells: The case of arrestins and GRKs. Crit Rev Biochem Mol Biol 50, 440–452. 10.3109/10409238.2015.1067185.

83. Gurevich, V.V., Gurevich, E.V., and Cleghorn, W.M. (2008). Arrestins as multi-functional signaling adaptors. Handb Exp Pharmacol, 15-37. 10.1007/978-3-540-72843-6_2.

84. Bychkov, E.R., Gurevich, V.V., Joyce, J.N., Benovic, J.L., and Gurevich, E.V. (2008). Arrestins and two receptor kinases are upregulated in Parkinson’s disease with dementia. Neurobiol Aging 29, 379–396. 10.1016/j.neurobiolaging.2006.10.012.

85. Gurevich, E.V., Gainetdinov, R.R., and Gurevich, V.V. (2016). G protein-coupled receptor kinases as regulators of dopamine receptor functions. Pharmacol Res 111, 1–16. 10.1016/j.phrs.2016.05.010.

86. Ahmed, M.R., Zheng, C., Dunning, J.L., Ahmed, M.S., Ge, C., Pair, F.S., Gurevich, V.V., and Gurevich, E.V. (2024). Arrestin-3-assisted activation of JNK3 mediates dopaminergic behavioral sensitization. Cell Rep Med, 101623. 10.1016/j.xcrm.2024.101623.

87. Ahmed, M.R., Bychkov, E., Gurevich, V.V., Benovic, J.L., and Gurevich, E.V. (2008). Altered expression and subcellular distribution of GRK subtypes in the dopamine-depleted rat basal ganglia is not normalized by l-DOPA treatment. J Neurochem 104, 1622–1636. 10.1111/j.1471-4159.2007.05104.x.

88. Lanza, K., Centner, A., Coyle, M., Del Priore, I., Manfredsson, F.P., and Bishop, C. (2021). Genetic suppression of the dopamine D3 receptor in striatal D1 cells reduces the development of L-DOPA-induced dyskinesia. Exp Neurol 336, 113534. 10.1016/j.expneurol.2020.113534.

89. Lanza, K., Meadows, S.M., Chambers, N.E., Nuss, E., Deak, M.M., Ferre, S., and Bishop, C. (2018). Behavioral and cellular dopamine D(1) and D(3) receptor-mediated synergy: Implications for L-DOPA-induced dyskinesia. Neuropharmacology 138, 304–314. 10.1016/j.neuropharm.2018.06.024.

90. Olson, M.F., and Schimmer, B.P. (1992). Heterologous desensitization of the human dopamine D1 receptor in Y1 adrenal cells and in a desensitization-resistant Y1 mutant. Mol Endocrinol 6, 1095–1102. 10.1210/mend.6.7.1354841.

91. Xu, W., Chen, C., Li, J.G., Dimattio, K., Wang, Y., Unterwald, E., and Liu-Chen, L.Y. (2013). PKA and ERK1/2 are involved in dopamine D(1) receptor-induced heterologous desensitization of the delta opioid receptor. Life Sci 92, 1101–1109. 10.1016/j.lfs.2013.04.006.

92. Misganaw, D. (2021). Heteromerization of dopaminergic receptors in the brain: Pharmacological implications. Pharmacol Res 170, 105600. 10.1016/j.phrs.2021.105600.

93. Pavon, N., Martin, A.B., Mendialdua, A., and Moratalla, R. (2006). ERK phosphorylation and FosB expression are associated with L-DOPA-induced dyskinesia in hemiparkinsonian mice. Biol Psychiatry 59, 64–74. 10.1016/j.biopsych.2005.05.044.

94. Ahmed, M.R., Bychkov, E., Kook, S., Zurkovsky, L., Dalby, K.N., and Gurevich, E.V. (2015). Overexpression of GRK6 rescues L-DOPA-induced signaling abnormalities in the dopamine-depleted striatum of hemiparkinsonian rats. Exp Neurol 266, 42–54. 10.1016/j.expneurol.2015.02.008.

95. Ahmed, M.R., Berthet, A., Bychkov, E., Porras, G., Li, Q., Bioulac, B.H., Carl, Y.T., Bloch, B., Kook, S., Aubert, I., et al. (2010). Lentiviral overexpression of GRK6 alleviates L-dopa-induced dyskinesia in experimental Parkinson’s disease. Sci Transl Med 2, 28ra28. 10.1126/scitranslmed.3000664.

96. Decressac, M., and Bjorklund, A. (2013). mTOR inhibition alleviates L-DOPA-induced dyskinesia in parkinsonian rats. J Parkinsons Dis 3, 13–17. 10.3233/JPD-120155.

97. Santini, E., Heiman, M., Greengard, P., Valjent, E., and Fisone, G. (2009). Inhibition of mTOR signaling in Parkinson’s disease prevents L-DOPA-induced dyskinesia. Sci Signal 2, ra36. 10.1126/scisignal.2000308.

